# Enforced dimerization between XBP1s and ATF6f enhances the protective effects of the unfolded protein response (UPR) in models of neurodegeneration

**DOI:** 10.1101/2020.11.17.387480

**Authors:** René L. Vidal, Denisse Sepulveda, Paulina Troncoso-Escudero, Paula Garcia-Huerta, Constanza Gonzalez, Lars Plate, Carolina Jerez, José Canovas, Claudia A. Rivera, Valentina Castillo, Marisol Cisternas, Sirley Leal, Alexis Martinez, Julia Grandjean, Hilal A. Lashuel, Alberto J.M. Martin, Veronica Latapiat, Soledad Matus, S. Pablo Sardi, R. Luke Wiseman, Claudio Hetz

**Author notes:** Corresponding author: Rene Vidal, Center for Integrative Biology, Universidad Mayor, Santiago, Chile. Claudio Hetz, Institute of Biomedical Sciences, Sector B, second floor, Faculty of Medicine, University of Chile, Independencia 1027, Santiago, Chile. P.O.Box 70086. or. www.hetzlab.cl.

## Abstract

Alteration to endoplasmic reticulum (ER) proteostasis is observed on a variety of neurodegenerative diseases associated with abnormal protein aggregation. Activation of the unfolded protein response (UPR) enables an adaptive reaction to recover ER proteostasis and cell function. The UPR is initiated by specialized stress sensors that engage gene expression programs through the concerted action of the transcription factors ATF4, ATF6f, and XBP1s. Although UPR signaling is generally studied as unique linear signaling branches, correlative evidence suggests that ATF6f and XBP1s may physically interact to regulate a subset of UPR-target genes. Here, we designed an ATF6f-XBP1s fusion protein termed UPRplus that behaves as a heterodimer in terms of its selective transcriptional activity. Cell-based studies demonstrated that UPRplus has stronger an effect in reducing the abnormal aggregation of mutant huntingtin and alpha-synuclein when compared to XBP1s or ATF6 alone. We developed a gene transfer approach to deliver UPRplus into the brain using adeno-associated viruses (AAVs) and demonstrated potent neuroprotection *in vivo* in preclinical models of Parkinson’s and Huntington’s disease. These results support the concept where directing UPR-mediated gene expression toward specific adaptive programs may serve as a possible strategy to optimize the beneficial effects of the pathway in different disease conditions.

## Introduction

The proteostasis network encompasses the dynamic integration of all cellular and molecular processes that ensure the proper folding and trafficking of proteins^1^. The endoplasmic reticulum (ER) is a central node of the proteostasis network, mediating the synthesis and quality control of near 30% of the total proteome. Different stress conditions can interfere with the function of the secretory pathway, leading to aberrant protein folding and resulting in a cellular state called ER stress^2, 3^. To recover ER proteostasis, cells activate an integrated signaling pathway known as the unfolded protein response (UPR), resulting in the establishment of adaptive outputs to decrease the extent of protein misfolding and enter into a new homeostatic state^4^. At the mechanistic level, activation of the UPR leads to the attenuation of protein translation, and the upregulation of multiple genes encoding for chaperones, foldases, and components of the protein quality control and degradation machinery (i.e. ER-associated degradation (ERAD) and autophagy)^5^. However, under sustained or irreversible ER stress, a terminal UPR results in cellular apoptosis^6, 7^. Overall, the UPR integrates stress signals toward the mitigation of ER stress or the induction of cell death, thus determining cell fate.

Three distinct ER-located stress sensors mediate the initiation of the UPR. These include inositol-requiring enzyme 1 (IRE1) alpha and beta, activating transcription factor-6 (ATF6) alpha and beta, and protein kinase RNA (PKR)-like ER kinase (PERK)^5^. IRE1α is a type-I Ser/Thr protein kinase and an endoribonuclease that upon activation catalyzes the processing of the mRNA encoding the transcription factor X-Box-binding Protein 1 (XBP1), excising a 26-nucleotide intron followed by a ligation reaction by RTCB^8^. This alternative splicing event shifts the coding reading frame of the mRNA to translate a stable and active basic leucine zipper (bZIP) transcription factor known as XBP1s^9–11^. XBP1s upregulates different genes involved in folding, ERAD, protein translocation into the ER, phospholipid synthesis among other components of the proteostasis network^12, 13^. The expression of XBP1s has an essential physiological role in sustaining the function of different organs and specialized secretory cells, including the brain, immune cells, liver, pancreas as well as other cell types and tissues (reviewed in^14^). Besides, IRE1α signals through the direct degradation of mRNAs and microRNAs by a process known as regulated IRE1-mediated decay (RIDD) ^15–17^, in addition to operating as a scaffold that binds adapter proteins and signaling molecules^18^. Activating transcription factor 6 alpha (ATF6α) is a type II transmembrane protein, that under ER stress is translocated to the Golgi apparatus where it is proteolytically processed, releasing the cytoplasmic fragment of ATF6α that contains a bZIP transcription factor (ATF6f, also known as ATF6p50)^19–21^. ATF6f upregulates genes implicated in ERAD and protein folding, in addition to modulate the expression of XBP1 mRNA^9, 22, 23^. Finally, PERK is a protein kinase that upon activation selectively inhibits the translation of proteins through the phosphorylation of eukaryotic translation initiator factor-2 (elF2α) at serine 51^24–26^. This event decreases the overload of misfolded proteins at the ER but also allows the selective translation of the mRNA encoding the transcription factor ATF4^27^. ATF4 regulates the expression of genes involved in redox control, amino acid synthesis, protein folding, and autophagy^28^, in addition to pro-apoptotic factors such as CHOP/GADD153 and members of the BCL-2 family of proteins^29^.

Chronic ER stress is emerging as a possible factor contributing to the pathogenesis of various human diseases including cancer, neurodegeneration, metabolic syndromes, inflammation, and fibrosis (reviewed in^4, 7, 30^). In fact, pharmacological targeting of the UPR has proven efficacy in various preclinical models of different diseases^31^. In this context, the ER proteostasis network is becoming an attractive target to treat diseases associated with protein misfolding and aggregation affecting the brain^32^. Signs of ER stress have been extensively reported in brain tissue derived from patients affected with Alzheimer’s disease (AD), Parkinson’s disease (PD), Huntington’s disease (HD), amyotrophic lateral sclerosis (ALS), frontotemporal dementia, and prion disease^33^. Genetic and pharmacological manipulation of distinct UPR components in preclinical models of neurodegeneration has demonstrated a complex scenario, where depending on the pathological condition and the specific signaling branch manipulated, the severity of the disease can be exacerbated or attenuated (reviewed in^34–36^). Since the transcriptional activity of both XBP1s and ATF6f is exclusively linked to the establishment of adaptive and pro-survival responses, gene therapy strategies to artificially enforce their expression have been exploited in the context of various neurodegenerative diseases^37^. For example, XBP1s overexpression reduces the aggregation and toxicity of disease-related proteins including mutant huntingtin (mHtt), tau, and amyloid β^38–41^. Also, XBP1s overexpression can improve neuronal function in models of AD at the level of synaptic plasticity^42^, enhance the survival of dopaminergic neurons against PD-inducing neurotoxins^43, 44^ or protect retinal ganglion cells from glaucoma^45^. The expression of XBP1s is also protective in models of mechanical injury to the spinal cord^46^ and peripheral nerves^47^. Moreover, XBP1s overexpression has been shown to reduce tissue damage in preclinical studies of heart failure ^48, 49^, liver diseases, and metabolic syndromes^50, 51^. ATF6 expression causes neuroprotection in early phases of experimental HD^52^, whereas activation of the pathway using small molecules can protect various tissues against ischemia-reperfusion including the brain^53–55^. Gene therapy to deliver active ATF6 also protects the heart against ischemia-reperfusion^56^. Overall, these selected studies support a concept where the improvement of ER proteostasis through the artificial enforcement of UPR gene expression programs improve cell function and survival in various disease settings.

Although the three main UPR signaling branches have been historically studied as individual entities, a few reports suggest the occurrence of signaling crosstalk at different levels. ATF6f and XBP1s are bZIP transcription factors that bind to related DNA CCAATN9CCACG cassettes^57, 58^. XBP1s and AFT6f have been suggested to functionally interact in cells suffering ER stress^9, 57, 59, 60^. Besides, both XBP1s and ATF6f share the control of a subpopulation of target genes, highlighting factors participating in ERAD and protein folding^13, 20^, in addition to the control of ER and Golgi biogenesis by enhancing phospholipid synthesis^61, 62^. Interestingly, the co-expression of ATF6f and XBP1s results in a particular remodeling of the ER proteostasis network that can be distinguished from the effects triggered by the single transcription factors^60^. Co-expression of XBP1s and ATF6f cooperates in the regulation of factors related to ER protein import (i.e. Sec11c), folding (i.e. PDIA10 and HYOU1), quality control (i.e. EDEM1 and DERL2), protein maturation (i.e. Sulf1), among others^63^. Because a previous study suggested that ATF6f and XBP1s may physically associate^22^, it was speculated that the formation of a heterodimer between the two transcription factors might translate into the establishment of divergent profiles of gene expression. This concept was also reinforced by other studies suggesting that XBP1, ATF6, and ATF4 can physically interact with other bZIP transcription factors, dictating the universe of target genes regulated (see examples in^51, 64–69^). However, the possible cooperative function of XBP1s and ATF6f in alleviating disease pathology has not been directly studied.

Here we developed a strategy to generate a fusion protein containing active XBP1s and ATF6f using a flexible linker to stabilize the formation of a heterodimer. This artificial protein is active and phenocopies the effects of co-expressing ATF6f and XBP1s in terms of further enhancing the expression of a specific set of genes. At the functional level, we found a strong anti-aggregation activity of UPRplus on mHtt and alpha-synuclein (α-synuclein) in cell culture models when compared with the expression of XBP1s or ATF6f alone. We developed a gene therapy approach to deliver XBP1s, ATF6f, or UPRplus into the mouse brain and compared their possible neuroprotective effects. Our results suggest that UPRplus has superior activity in providing neuroprotection and reducing the aggregation of disease-related proteins *in vivo.* Thus, UPR transcription factors cooperate to regulate gene expression and restore proteostasis in cells undergoing protein misfolding.

## Results

### Design of an active ATF6f/XBP1s fusion protein

To study the possible biological function of the XBP1s/ATF6f heterodimer, we generated different constructs that may stabilize the intramolecular formation of a heterodimer. We fused XBP1s and ATF6f with three different linkers (L4H4, LF, and LFG) that provide flexibility between the two transcription factors^70, 71^. We included hydrophilic helix-peptide linkers (GGGGS)n, previously described to force the interaction between two fluorescent proteins^72^. These linkers vary in their amino acid composition and length required to confer the necessary flexibility for DNA binding. We generated six variants of the fusion proteins in two different ways, using ATF6f in the amino or carboxyl-terminal positions (Figure 1A), in addition to a hemagglutinin (HA) epitope in the carboxyl terminus for detection. These constructs were expressed in HEK293T cells to measure their subcellular localization and stability. We determined the steady-state expression of these constructs and observed that the fusion proteins containing ATF6f at the N-terminal region showed reduced expression levels whereas the location of XBP1s at the N-terminus was associated with increased populations of truncated fragments (Figure 1B). Importantly, all six constructs were preferentially localized inside the nucleus, consistent with the expression of the active forms of XBP1s and ATF6f (Figure 1C). Next, we compared the transcriptional activity of these fusion proteins using a luciferase promoter reporter controlled by a UPR-responding element (UPRE)^13^. Although all six constructs were active, the fusion construct with ATF6f in the N-terminal region presented higher transcriptional activity despite lower expression levels (Figure 1D, green bars). As a control, XBP1s and ATF6f were co-expressed, showing similar activity to the fusion constructs containing ATF6f at the N-terminal region (Figure 1D, black bar).

**Figure 1.**
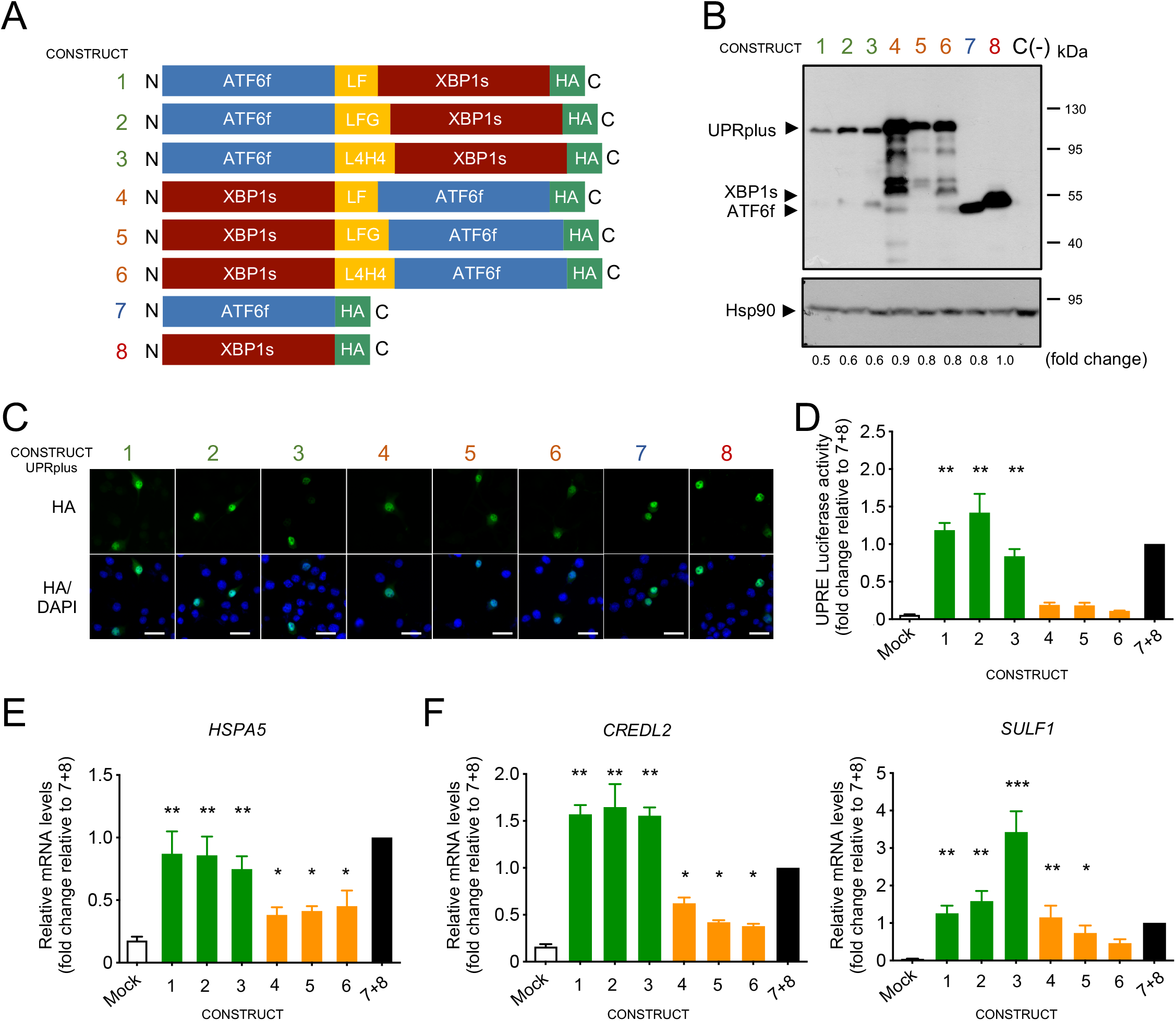
Generation of ATF6f and XBP1s fusion constructs. (A) Diagram of AAV constructs generated to deliver active UPR transcription factors. Different artificial heterodimers were generated by fusing ATF6f and XBP1 s using 3 different linker sequences (yellow boxes) by combining their positions in the C-terminal and N-terminal regions. All constructs contain an HA tag at the C terminal region (green box) for the detection of the expression of the transgene. (B) HEK293T cells were transiently transfected with the 6 fusion constructs described in panel *A,* in addition to XBP1s-HA or ATF6f-HA alone and empty vector C (-). After 48 h of expression, cell extracts were analyzed by western blot using an anti-HA antibody. Hsp90 was monitored as a loading control. Fold change are showed related to XBP1s expression levels (C) In parallel, cells described in *B* were analyzed by immunofluorescence using an anti-HA antibody (green). Co-staining with the nuclear marker Hoechst (blue) was performed. Scale bar 10 μm. (D) HEK293T cells were transiently co-transfected with the 6 variants of pAAV-UPRplus or single constructs together with the UPRE-luciferase reporter and renilla constructs. After 48 h luciferase activity was measured using a luminometer. (E) HEK293 cells were transiently transfected with the 6 versions of UPRplus or control vectors. After 48 h, the mRNA levels of the indicated UPR-target gene were measured by real-time RT-PCR. All samples were normalized to β-*actin* levels. mRNA levels are expressed as fold increase over the value obtained in control cells transfected with an equivalent 1:1 mixture of individual XBP1s and ATF6f expression vectors. In D and E, the mean and standard error is presented of three independent experiments. Statistical analysis was performed using Dunnett’s multiple comparisons test (*: *p* < 0.05; **: *p* < 0.01; ***: *p* < 0.001).

Then, we measured the relative mRNA levels of the canonical UPR target gene *HSPA5* (also known as *BiP/GRp78)* in HEK293T cells transiently expressing all six constructs. The presence of ATF6f in the N-terminal region showed a greater induction of HSPA5 (Figure 1E, green bars) compared to the constructs containing XBP1s at the N-terminus (Figure 1E, orange bars). Again, the induction levels of HSPA5 by the first three constructs were similar to the one generated by the co-expression of ATF6f and XBP1s (Figure 1E, black bar). A previous study indicated that the simultaneous overexpression of XBP1s and ATF6 resulted in a stronger upregulation of a small subset of UPR-target genes, including *CRELD2* and *SULF1*^60^. In agreement with those observations, expression of the fusion constructs containing ATF6f in the N-terminal region showed a greater induction of the of *CREDL2* and *SULF1* mRNAs (Figure 1F, green bars), where the third variant had a greater effect in increasing *SULF1* levels. We also controlled the activity of single constructs by measuring the upregulation of ERdj4 (XBP1s target gene) and Grp94 (ATF6f target gene (Figure S1A).

Finally, we performed disorder predictions to determine the flexibility of all six constructs using ESpritz, a software designed to determine amino acids with missing atoms in X-ray solved structures, associated with mobile amino acids within protein structures^73^. This analysis predicted the lowest flexibility of the L4H4 linker in the construct ATF6f-L4H4-XBP1s-HA than the other constructs (Figure S1B). Based on these results, the stability of the fusion proteins, their subcellular localization, and transcriptional activity, we selected the ATF6f-L4H4-XBP1s fusion protein for further studies, a construct hereon termed UPRplus.

To compare the activity of UPRplus with the co-expression of XBP1s and ATF6f, we employed the original experimental system used to identify the functional interaction between both transcription factors. In that setting, HEK-Rex cells were engineered to express XBP1s under the control of doxycycline (DOX), whereas ATF6f was constrictively expressed with a destabilization domain that induces its degradation and can be rescued with the addition of the pharmacological chaperone trimethoprim (TMP). As previously reported, the administration of DOX and TMP to these cells synergized to upregulate the expression of *SULF1* when compared with single treatments (Figure S1C). We then compared the effects of expressing UPRplus, XBP1s, ATF6f or the co-expression of both transcription factors using the same cells as isogenic background. Remarkably, the expression of UPRplus have a stronger capacity to upregulate endogenous *SULF1* when compared with the co-expression of XBP1s and ATF6f (Figure S1C).

### Interactions within the XBP1s and ATF6f sequences are necessary for gene expression control

Both XBP1s and ATF6f belong to the basic leucine zipper (bZIP) transcription factor family, which are known to bind DNA through the formation of homodimers or heterodimers with other members of their family^9, 21, 74–76^. We hypothesized that UPRplus may bind promoter regions either by intramolecular interactions that result in the binding of XBP1s and ATF6f sequences within the single fusion constructs. However, UPRplus may also bind endogenous partners (i.e. XBP1s, ATF6f, or others) or could have homotypic interactions between individual UPRplus fusion proteins. To detect the size of the DNA-protein complex generated by UPRplus, we performed an electrophoretic mobility shift assay (EMSA) using purified nuclear extracts from HEK293T cells transfected with XBP1s, ATF6f, or UPRplus expression vectors. After 48 h, we detected an enrichment of the proteins in the nuclear fraction (Figure S2A). We incubated the nuclear extracts with a conserved DNA sequence containing a UPRE cassette. We observed a clear shift in the migration pattern of the UPRE probe when ATF6f or XBP1s was present (Figure 2A, lanes 2 and 5). As a control, we performed competition experiments using a non-labeled DNA probe, which resulted in a reduction in the amount of the DNA-protein complexes formed by XBP1 (Figure 2A, lane 3), while ATF6 and UPRplus complex was slightly modified (Figure 2A, lanes 6 and 9). Importantly, the signal was absent when a UPRE mutant probe that lacks the minimal recognition region for ATF6f and XBP1s was used (Figure 2A, lanes 4 and 7). Interestingly, a DNA-protein complex of similar size to the one detected when XBP1s or ATF6f were expressed was detected after expressing UPRplus (Figure 2A, lane 8; see controls line 9 and 10). These results support the idea that UPRplus might bind to DNA as a single molecule.

**Figure 2.**
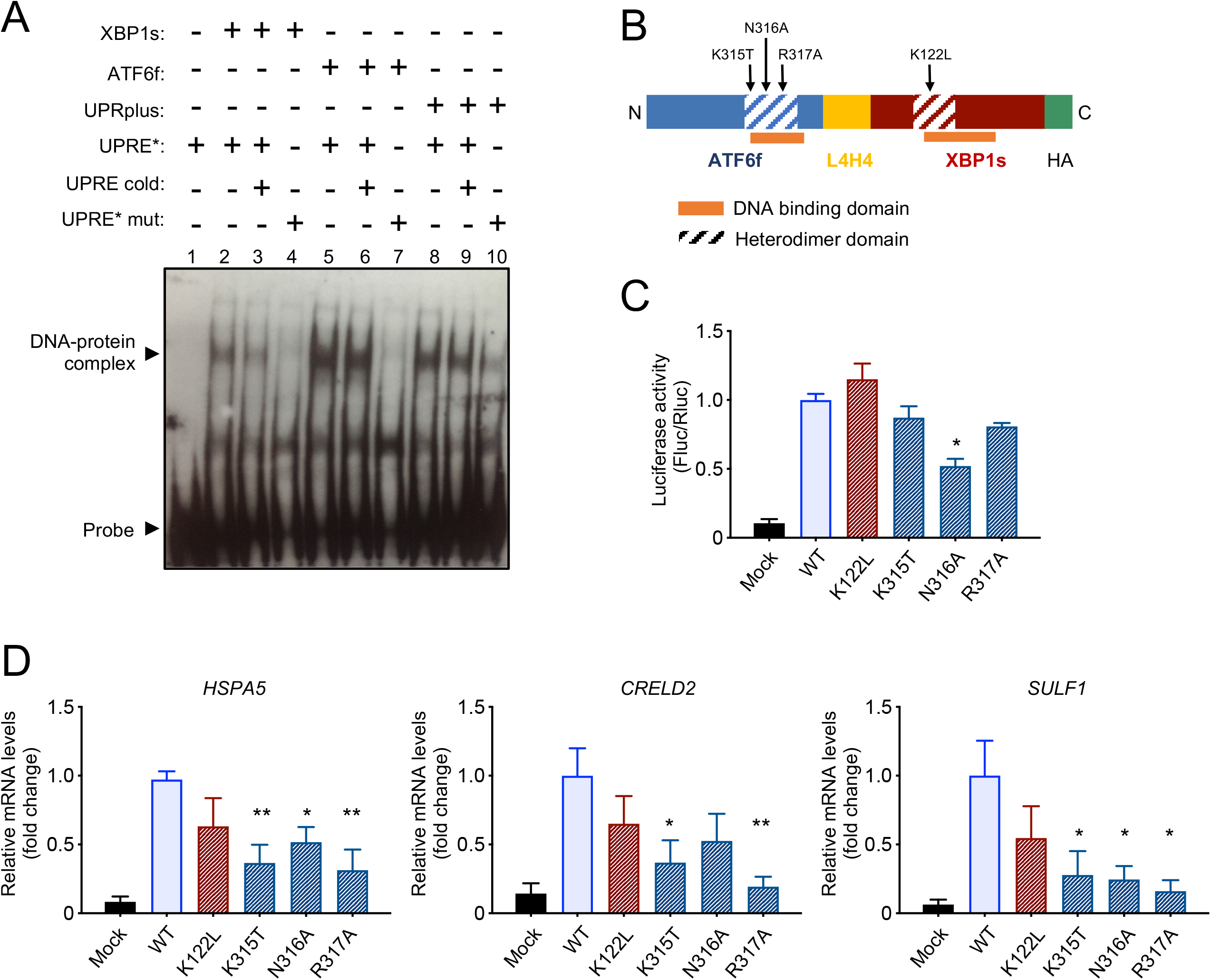
UPRplus binds to a UPR response element and requires the dimerization interface for gene expression regulation. (A) Nuclear extracts obtained from HEK293T cells transfected with pAAV-XBP1s-HA (2-4 lanes), pAAV-ATF6f-HA (5-7 lanes), pAAV-UPRplus-HA (8-10 lanes) or empty vector were incubated with labeled UPRE* probe. As a control, the competition was performed with unlabeled (cold probe, lanes 3, 6 or, 9) or mutated probes (UPRE* mut, line 7, 4, or 10). The asterisk represents the labeled probe. The DNA-protein complexes were separated on non-denaturing polyacrylamide gels and analyzed by western blot using an anti-biotin antibody. (B) Schematic representation of the UPRplus construct is indicated, highlighting the heterodimer domains of both ATF6f and XBP1s (dashed boxes). Point mutations generated in these domains are indicated. Yellow box: linker region. Green box: HA tag. (C) HEK293T cells were transiently co-transfected with UPRplus (UPRplus WT), or four single mutant versions of (K122L, K315T, N316A, and R317A), in addition to empty vector (Mock) together with the UPRE-luciferase reporter and renilla constructs. After 48 h, luciferase activity was measured using a luminometer. (D) HEK293T cells were transiently transfected with DNA constructs described in *C* and after 24 h, *HSPA5, CRELD2,* and *SULF1* mRNA levels measured by real-time RT-PCR. All samples were normalized to β*-*actin levels. mRNA levels are presented as fold increase over the value obtained in control cells transfected with the UPRplus WT version. In *C* and *D*, the mean and standard error is presented of three independent experiments. Statistical analysis was performed using Dunnett’s multiple comparisons test (*: *p* < 0.05; **: *p* < 0.01).

Several studies have mapped the domains required for DNA binding^77–79^, in addition to the possible theoretical regions needed for homotypic dimerization of bZIP transcription factors^80, 81^. These domains are overlapped in the primary structure of ATF6f and XBP1s (Figure 2B). To inactivate one of the two transcription factors contained in UPRplus, we generated specific point mutations in the dimerization domains of ATF6f and XBP1s present in the UPRplus sequence to then evaluate the effects in their transcriptional activity (Figures 2B and S2B). We mutated conserved single polar amino acid residues by replacing them by uncharged residues (K122L for XBP1s and K315T, N316A, or R317A for ATF6f) (Figures S2B and S2C). A homology model using as a template the 5T01 coordinates (human c-Jun bound to DNA) predicted that K315T, N316A, and R317A mutations are located at the DNA binding region of ATF6, together with the K122L mutation at the DNA binding region of XBP1s (Figure S2D).

Remarkably, a 50% reduction in the UPRE-luciferase activity was observed when the N316A mutation was introduced to UPRplus (Figure 2C; see the expression of mutants in Figure S2E and S2F). Furthermore, the K315T, N316A, and R317A mutations also diminished the transcriptional activity of UPRplus when the mRNA levels of HSPA5, CREDL2, and SULF1 were measured using real-time RT-PCR (Figure 2D). These results further support the idea that UPRplus regulate gene expression through intramolecular dimerization of the XBP1 s and ATF6f domains within the fusion protein.

### UPRplus expression reduces abnormal protein aggregation in cell-based models

Accumulating evidence support a functional role of the UPR in reducing abnormal protein aggregation in different neurodegenerative diseases^7, 33, 34, 82^. Thus, we decided to compare the anti-aggregation activity of UPRplus with XBP1s and ATF6f in cell culture models of proteinopathies. We first expressed a peptide containing 79 polyglutamine fused to EGFP (polyQ_79_-EGFP) in the neuroblastoma cell line Neuro2A. We transiently coexpressed polyQ_79_-EGFP together with UPRplus, XBP1s, ATF6f, or control vector, and then examine the aggregation levels by western blot and filter trap. A decrease in the accumulation of high molecular weight (HMW) and detergent-insoluble polyQ_79_-EGFP species was detected when all three constructs were tested using both assays (Figures 3A-3D). Importantly, the inhibitory effects of UPRplus in the accumulation of polyQ_79_-EGFP aggregates were more potent than ATF6f or XBP1s alone (Figures 3A-3D and S6A). Virtually identical results were obtained when we quantified the presence of intracellular polyQ_79_-EGFP inclusions using fluorescent microscopy (Figure 3E). We then compared the efficacy of UPRplus in reducing polyQ_79_-EGFP aggregation with the co-expression of XBP1s and ATF6f using the DOX-TMP inducible system (Figure S1D). Remarkably, UPRplus expression was more efficient in reducing abnormal protein aggregation when compared with the co-expression system (Figure S1D).

**Figure 3.**
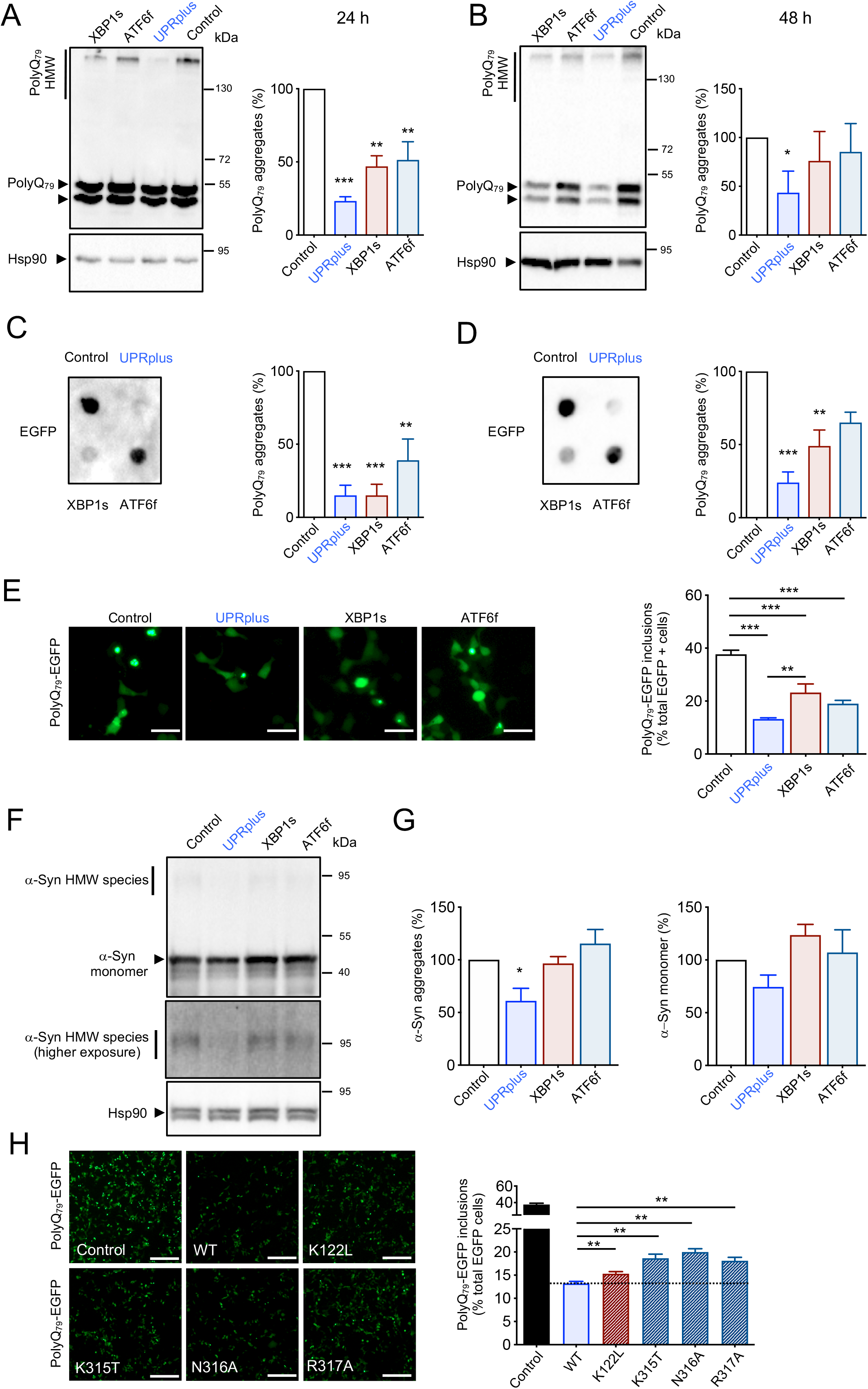
UPRplus expression reduces mutant huntingtin and α-synuclein aggregation. (A-B) Neuro2A cells were transiently co-transfected with expression vectors for polyQ_79_-EGFP and XBP1s, ATF6f, UPRplus, or empty vector (Control). After 24 (A) or 48 h (B), polyQ_79_-EGFP detergent-insoluble aggregates were measured in cell extracts prepared in Triton X100 by western blot. Levels of Hsp90 were measured as the loading control. Left panel: high molecular weight (HMW) polyQ_79_-EGFP aggregates were quantified. (C-D) polyQ_79_-EGFP detergent-insoluble aggregates were measured by filter trap assay after 24 (C) or 48 h (D) of transfection (right panel). Left panels: polyQ_79_-EGFP aggregates were quantified. (E) In cells described in *A*, polyQ_79_-EGFP intracellular inclusions were quantified after 48 h of expression by fluorescent microscopy (right panel). Scale bar, 20 μm. Left panel: The number of cells displaying intracellular inclusions was quantified in a total of at least 300 cells per experiment. (F) HEK293T cells were transiently co-transfected with expression vectors for α-synuclein-RFP (α-syn) together with expression vectors for UPRplus, ATF6f, XBP1s, or empty vector (Control). After 48 h, α-syn aggregates were measured in cell extracts prepared in 1% Triton X-100 by western blot (upper panel). Middle panel: higher exposure of the upper panel highlighting HMW species of α-syn. Levels of Hsp90 were monitored as the loading control (lower panel). (G) α-synuclein-RFP HMW species (left panel) and α-syn monomers (right panel) were quantified. (H) Neuro2A cells were transiently co-transfected with expression vectors for polyQ_79_-EGFP together with empty vector (control), UPRplus WT, or the UPRplus mutants K122L, K315T, N316A, and R317A. After 48 h, the accumulation of polyQ_79_-EGFP intracellular inclusions was visualized by fluorescence microscopy (right panel). Scale bar, 100 μm. Left panel: The number of cells displaying intracellular inclusions was quantified in a total of at least 300 cells per experiment. In A-G, the mean and standard error is presented of three independent experiments. In G four independent experiments were quantified. Statistical analysis was performed using Dunnett’s multiple comparisons test (*: *p* < 0.05; **: *p* < 0.01).

We then evaluated the possible effects of UPRplus on the aggregation of other aggregation-prone proteins. We co-expressed human α-synuclein together with UPRplus or control vectors in HEK293T cells and examined the aggregation levels by western blot. UPRplus expression induced a significative decrease of α-synuclein aggregation levels, in addition to a tendency in reducing its monomeric form (Figures 3F and 3G). Unexpectedly, the expression of XBP1s or ATF6f alone did not modify the levels of α-synuclein aggregation (Figures 3F and 3G).

We then explored the consequences of mutating the DNA binging interphase of UPRplus on the aggregation of polyQ_79_-EGFP. A slight but significant reduction in the accumulation of polyQ_79_-EGFP inclusions was observed while the different point mutants of UPRplus were tested (Figures 3H and S3A). Similar results were obtained when protein aggregation of polyQ_79_-EGFP was determined by western blot (Figure S3B). Overall, our results suggest that the expression of the UPRplus fusion construct has a stronger biological activity in reducing abnormal protein aggregation when compared to single XBP1s or ATF6 molecules.

### UPRplus remodels the proteostasis network toward improving protein folding

To assess global changes in gene expression triggered by UPRplus, we performed quantitative proteomics using tandem mass tags for relative protein quantification and Multi-Dimensional Protein Identification Technology (MuDPIT)^83^. We transfected UPRplus, XBP1s, ATF6f constructs or empty vector in HEK293T cells followed by proteomic analysis after 48 h. The expression of a total of 123 proteins was identified to be modulated showing a *q* value < 0.15 and a minimum log2 > 0.1-fold change (Tables S1 and S2). We observed a low correlation between the changes in the proteome triggered by UPRplus with the ones induced by ATF6 (R2 = 0.51), highlighting the upregulation of several factors involved in protein folding (Figure 4A). We also correlated the proteome changes induced by UPRplus and XBP1s, observing even a lower correlation compared with ATF6 (R2 = 0.31), suggesting the induction of distinct patterns of gene expression.

**Figure 4.**
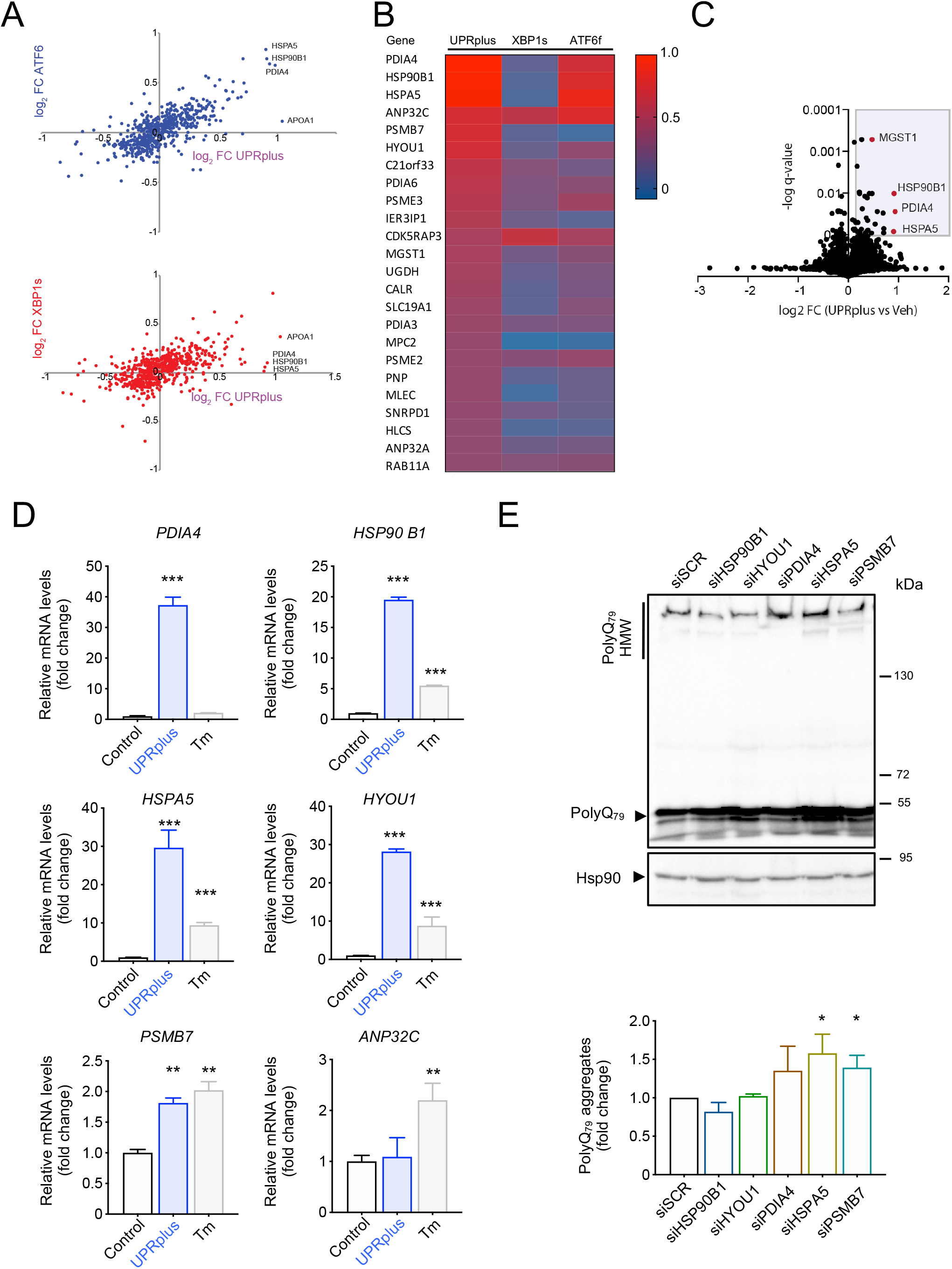
UPRplus expression remodels proteostasis pathways. (A) Quantitative proteomics was performed in protein extracts from HEK293T cells infected for 48 h with the following viral particles: AAV-UPRplus, AAV-XBP1s, AAV-ATF6f, or AAV-empty (vehicle). The data was analyzed and plotted of log2 fold change (FC) for proteins identified in MuDPIT analysis. Plot showing the correlation between gene expression in cells expressing UPRplus or ATF6f (upper panel) and UPRplus or XBP1s (bottom panel). Only genes whose expression is significantly affected (FDR < 0.01) are shown. (B) Heat map analysis showing differential protein expression patterns in ATF6f, XBP1s, or UPRplus overexpression conditions. Color from red to blue indicates high to low expression. (C) Volcano plot showing the correlation between protein expression of HEK293T cells infected with AAV-UPRplus versus AAV-empty (vehicle) viral particles. Only genes whose expression is significantly affected (FDR < 0.01) are shown. (D) The mRNA levels of selected UPR-upregulated genes were monitored in HEK293T cells infected with AAV-empty (control) or AAV-UPRplus viral particles. After 48 h the relative mRNA levels of the indicated genes were measured by realtime RT-PCR. As a positive control, cells were treated with 1 μg/ml tunicamycin for 8 h (Tm). All samples were normalized to β*-actin* levels. mRNA levels are expressed as fold increase over the value obtained in the control condition. (E) HEK293T cells were transiently transfected with siRNA against the 6-top gene genes upregulated by UPRplus. A scrambled siRNA (siScr) was used as control. 24 h later cells were transfected with a polyQ_79_-EGFP expression vector followed by western blot analysis after 24 h of expression. Levels of Hsp90 were analyzed as a loading control. Bottom panel: polyQ_79_-EGFP high molecular weight (HMW) species were quantified. In panels D-E, the mean and standard error is presented of three independent experiments. Statistical analysis was performed using Dunnett’s multiple comparisons test (*: *p* < 0.05; **: *p* < 0.01; ***: *p* < 0.001).

We then determined the top genes upregulated by UPRplus (Figure 4B and 4C) that may operate as effectors of its anti-aggregation activity. We selected the first six top hits for further validation and determine the mRNA levels using real-time RT-PCR. From these experiments, we were able to confirm a strong upregulation of *PDIA4* (a protein disulfide isomerase known as ERp72), *HSP90B1* (also known as GRP94), *HSPA5* (known as BiP), *PSMB7* (proteasome 20S subunit beta 7) and *HYOU1* (known as GRp170, ORP-150 or HSP12A), whereas the mRNA levels of *ANP32C* (an Hsp90 client, also known as pp32r1) were not altered (Figure 4D). As a control, cells were also stimulated with the ER stressor tunicamycin to confirm their regulation by ER stress (Figure 4D). Analysis of the proximal promoter regions of these 5 selected genes indicated the presence of canonical ER stress-response element (ERSE) I (CCAAT-N9-CCAC) and ERSE II (ATTGG-N1-CCACG) on a range of 15 kb upstream of the transcriptional starting site (TSS)^84, 85^ (Figure S4A). Moreover, we determined the transcription factor binding motifs to XBP1 s and ATF6f present in the promoter region of the five selected genes using the FIMO tool and the CIS-BP repository and found several sites for both XBP1 s and ATF6f on different positions upstream of the TSS. We identified putative binding sites where ATF6f, XBP1s, or both could bind to the promoter regions for each gene (Figures S4B-S4D).

To define the possible contribution of the genes identified to be regulated by UPRplus at the functional level, we performed knockdown experiments using a pool of small interfering RNAs (siRNAs) and confirmed their efficacy using HEK293T cells (Figure S5A). Next, we transfected the different pool of siRNAs followed by the expression of polyQ_79_-EGFP to measure the aggregation levels. A significant increase in the accumulation of polyQ_79_-EGFP aggregation was observed when HSPA5 (BiP) or PSMB7 were knocked down (Figure 4E). To control transfection efficiency, we monitored the mRNA levels of EGFP (Figure S5B). These results suggest that the upregulation of the ER chaperone BiP and a proteasome subunit PSMB7 may contribute to improve proteostasis and reduce abnormal protein aggregation in cells expressing UPRplus.

### UPRplus reduces mutant huntingtin aggregation *in vivo*

Gene therapy strategies to deliver the active forms of ATF6f or XBP1s have proven to be beneficial in various disease models (reviewed in^37^). To test the therapeutic potential of UPRplus *in vivo* in the context of neurodegenerative diseases, UPRplus, XBP1s, ATF6f, or empty vector were packed into adeno-associated virus (AAVs) using serotype 2 to transduce neurons *in vivo.* We validated the activity of these viral particles using primary cortical neurons, by adding the viral particles at 1 day of culture and then monitor the expression of UPR target genes. We observed a strong upregulation of *HspA5, HerpUD1, Credl2, and Hyou1* mRNA levels in neurons expressing UPRplus (Figure 5A). In sharp contrast, expression of XBP1s or ATF6f alone resulted in poor induction in primary neuronal cultures when compared to UPRplus (Figure 5A; see controls in Figure S6B).

**Figure 5.**
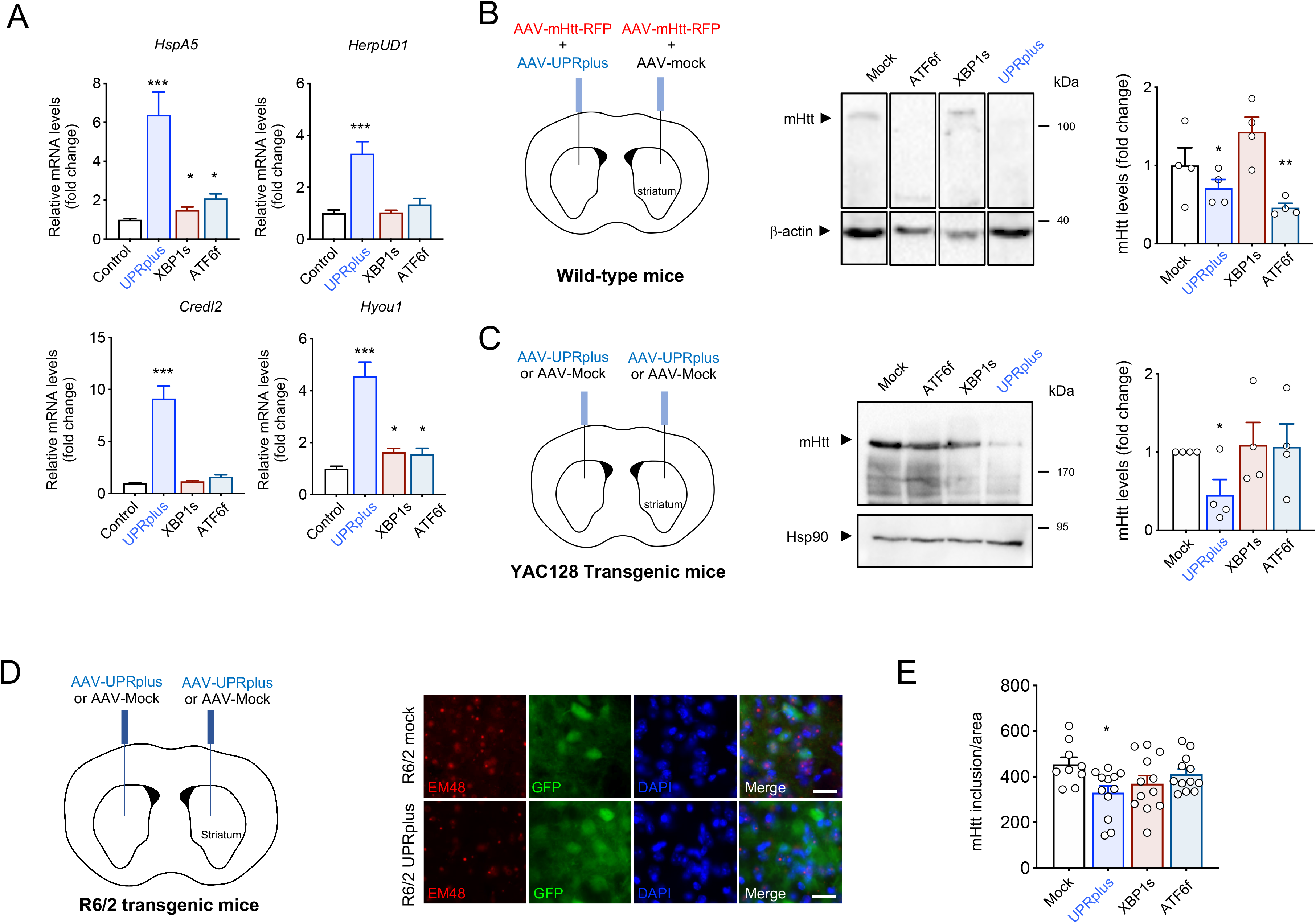
UPRplus expression decreases mutant huntingtin aggregation *in vivo*. (A) Primary cortical neurons were infected at 1 day in vitro (DIV) with adeno-associated virus (AAV) encoding for UPRplus, ATF6f, XBP1s, or empty vector (control). After 6 DIV, expression levels of UPR-target genes were measured by real-time RT-PCR. All samples were normalized to β*-actin* levels. mRNA levels are expressed as fold increase over the value obtained in control cells infected with AAV-empty (control). (B) Three-months old wild type mice were co-injected into the striatum by stereotaxis with a mixture of AAVs encoding a mHtt construct (Htt588^Q95^-mRFP) together with AAV-UPRplus, AAV-XBP1s, AAV-ATF6f, or AAV-Mock (control). Schematic representation of the experimental strategy is shown (left panel). Animals were then euthanized 2 weeks post-injection and brain striatum tissue was dissected for western blot analysis using an anti-polyQ antibody. β-actin levels were analyzed as a loading control (middle panel). High molecular weight (HMW) mHtt aggregates were quantified and normalized to β-actin levels (right panel) (AAV-Mock: n = 4; AAV-UPRplus: n = 4; AAV-XBP1s: n = 4; AAV-ATF6f: n = 4). (C) Three-month-old YAC128 mice were injected into the striatum with AAV-UPRplus, AAV-ATF6f, AAV-XBP1s, or AAV-Mock vector (control) using bilateral stereotaxis surgery. Schematic representation of the experimental strategy is shown (left panel). Four weeks later the striatum region was dissected and mHtt aggregation levels analyzed by western blot using an anti-polyQ antibody (middle panel). mHtt aggregates levels were quantified and normalized to Hsp90 levels (right panel). In B and C, the mean and standard error is presented for the analysis of four animals per group. (D) R6/2 mice were injected at 4 weeks of age with a mixture of AAV-EGFP and AAV-XBP1s (n = 4), AAV-ATF6f (n = 4), AAV-UPRplus ( n = 4) or AAV-Mock (control) ( n = 3) into the striatum using bilateral stereotaxis (left panel). Four weeks after injection, the brain was extracted and coronal slices from the striatum were obtained. mHtt was detected using the anti-huntingtin EM48 antibody (red) by fluorescence microscopy (red). EGFP expression was monitored as control for the injection (green). Nuclei were stained using DAPI (blue) (Scale bar: 20 μm) (right panel). (E) High-resolution images of the slices were obtained and quantification of mHtt was performed using Image J software. The quantification of the number of mHtt inclusions was performed by total area. Statistical analysis was performed using Dunnett’s multiple comparisons test (*: *p* < 0.05; **: *p* < 0.01).

To test the efficacy of UPRplus on reducing protein aggregation *in vivo,* we delivered the constructs to a viral mouse model of Huntington’s disease based on the expression of a mHtt fragment correspond to the first 588 amino acids that contain a track of 95 glutamines fused to monomeric RFP^38, 86^. This construct was delivered to the striatum using AAVs serotype 2 (termed AAV-Htt588^Q95^-mRFP). We co-injected AAV-Htt588^Q95^-mRFP particles together with AAV-UPRplus, AAV-ATF6f, AAV-XBP1s, or empty vector into the right striatum of adult mice (3-months old) using stereotaxis. After 3 weeks post injection, the expression of UPRplus resulted in a significant decrease in Htt588^Q95^-mRFP aggregation levels assessed using western blot from dissected striatal tissue (Figures 5B and S10). In this setting, the local expression of ATF6f also resulted in a similar reduction of mHtt aggregation (Figures 5B and S10). Unexpectedly, XBP1s expression did not have clear effects under these experimental conditions (Figures 5B and S10), which might be due to the use of different titers and AAV purification systems, in addition to differences in the DNA constructs (i.e. human versus mouse cDNA, the use of HA tag and the presence of a GFP cassette).

We moved forward and tested the possible effects of UPRplus on the mHtt levels using transgenic mice that express the full-length protein containing a track of 128 glutamines including the endogenous promoter on an yeast artificial chromosome (known as YAC128^87^). We performed bilateral stereotaxic injection of AAV-UPRplus, AAV-ATF6f, AAV-XBP1s, or empty vector (control) into the striatum of 3-month old YAC128 mice. Four months after the injection, animals were euthanized, and the striatal tissue was dissected for biochemical analysis using an anti-polyQ antibody that only recognizes mutant but not wild type Htt. We confirmed the expression of all transgenes in striatal tissue derived from YAC128 injected mice (Figure S6C). Remarkably, the expression of UPRplus led to a 50% reduction in the levels of full-length mHtt (Figure 5C). Unexpectedly, with the viral titers and time point used in this study, the delivery of ATF6f or XBP1s into the brain of YAC128 mice did not alter mHtt levels (Figure 5C).

To complement our studies, we validated our results on a third animal model of HD, the R6/2 mice, a transgenic line that expresses exon 1 of human huntingtin containing ~150 CAG repeats^88^ which allows the visualization of intracellular mHtt inclusions. 4 weeks old mice were bilaterally injected with AAV-XBP1s, AAV-ATF6f, AAV-UPRplus or AAV-empty into the striatum, followed by tissue immunofluorescence analysis of the brains six weeks later (Figure 5D). We observed a significant reduction in the number of mHtt-positive inclusions in mice injected with AAV-UPRplus (Figure 5D). Quantification of these experiments indicated a reduction of near 40% in the brain of R6/2 mice treated with AAV-UPRplus, whereas the expression of either XBP1s or ATF6f alone did not have a significant effect (Figure 5E). Taken together, these results validate the activity in reducing aggregation-prone protein of UPRplus *in vivo* using three different models of proteinopathies.

### UPRplus protects dopaminergic neurons in a pharmacological model of Parkinson’s disease

ER stress has been suggested as a relevant factor mediating dopaminergic neuron loss in various models of Parkinson’s disease (PD) (reviewed in^89^), including the use of pharmacological agents that mimics PD features in mice (see examples in^90–93^). At the functional level, previous studies indicated that the local injection of recombinant viruses to express XBP1s into the substantia nigra pars compacta (SNpc) reduces the degeneration of dopaminergic neurons triggered by PD-inducing neurotoxins^43, 44^, whereas ATF6 deficiency exacerbate the rate of neuronal loss^94, 95^. Thus, we moved forward and evaluated the potential therapeutic effects of UPRplus in a model of PD *in vivo.* We injected AAV-UPRplus, AAV-XBP1s, AAV-ATF6f, or empty vector into the SNpc by stereotaxis. After 14 days of expression, neuronal degeneration was induced by the unilateral injection of 6-hydroxydopamine (6-OHDA) into the striatum followed by histological analysis one week later (see schema in Figure 6A). We confirmed the expression of UPRplus in the SNpc using an anti-HA antibody (Figure 6B), in addition to the upregulation of BiP in dopaminergic neurons (Figure 6C).

**Figure 6.**
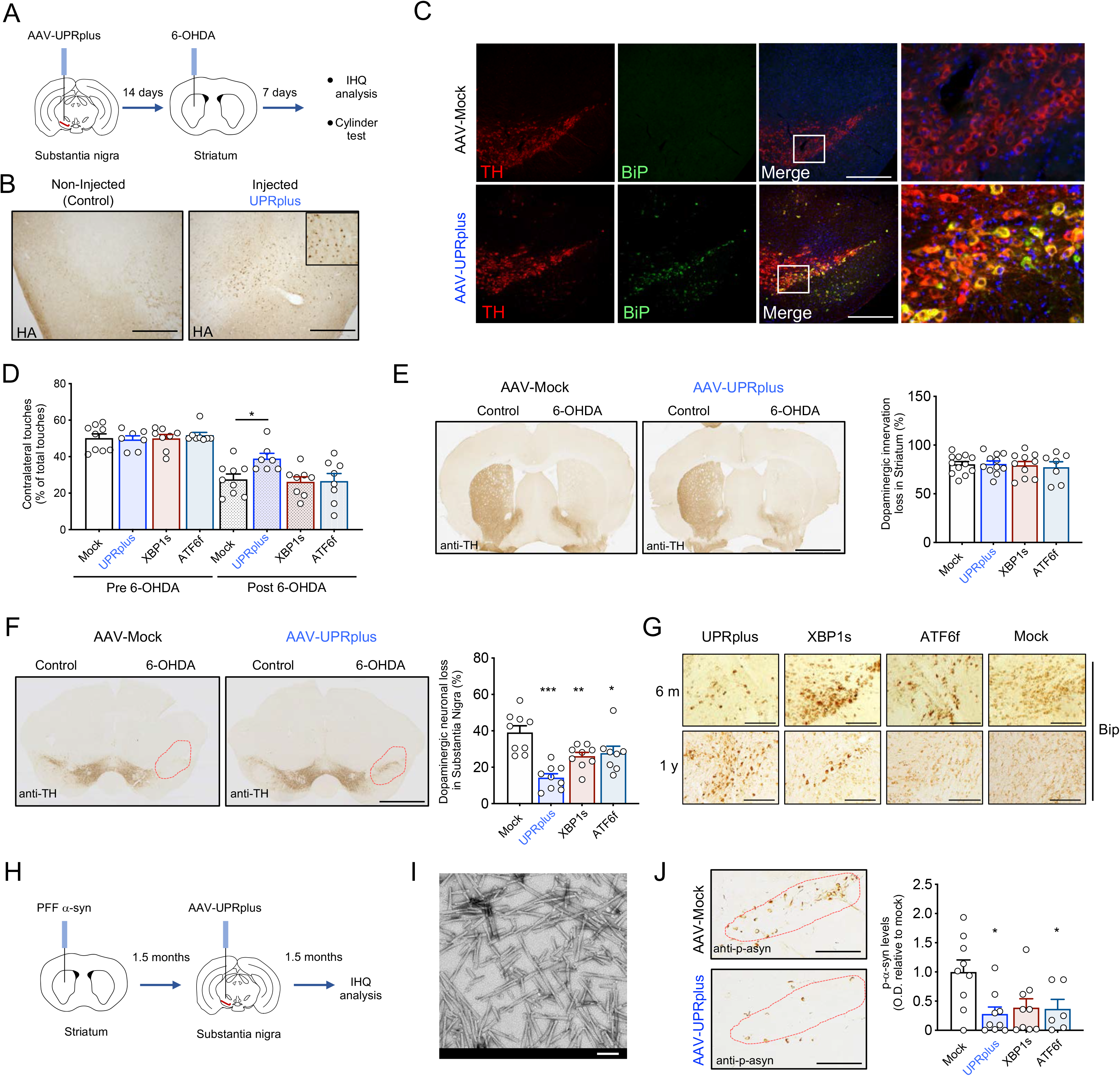
UPRplus protects dopaminergic neurons in preclinical model of PD. (A) Experimental design to evaluate the effects of UPRplus in a pharmacological PD model. Animals were injected with AAV particles expressing UPRplus or control vectors into the substantia nigra pars compacta (SNpc) using brain stereotaxis. Two weeks later, animals were exposed to 6-OHDA into the striatum followed by behavioral and histological analysis one week later. (B) The expression of UPRplus was monitored in the brain using immunohistochemistry with an anti-HA antibody (Scale bar: 200 μm). (C) Immunofluorescence analysis of tyrosine hydroxylase (TH; red) and BiP (green) was performed in brain tissue derived from AAV-mock (upper) or AAV-UPRplus (bottom) injected animals. Hoechst staining of the nucleus was also performed. The third panel shows merged images of the three staining (Scale bar: 200 μm). Right panel shows high magnification of the white square region of merged images. (D) Three-months old wild type mice were injected with AAV-UPRplus AAV-ATF6f, AAV-XBP1s, or AAV-mock (control) into the SNpc and then exposed to 6-OHDA as described in (A). Quantification of the percentage of contralateral touches relative to total touches (both forepaws) obtained before and 1 week after the injection of 6-OHDA (pre and post-6OHDA) is indicated. Data represent the mean and standard error of 6-8 animals per group. (E) Immunohistochemistry analysis was performed in striatal sections from animals described in (D) to quantify 6-OHDA–induced denervation in both injected (6-OHDA) and non-injected (control) hemispheres (scale bar: 1 mm) (left panel). The integrated density of pixel intensity was calculated from images of anti-TH immunohistochemistry covering the entire striatum and expressed as the percentage of TH loss relative to the control side. Data represent the mean and standard error from 6-8 animals per group (right panel). (F) Immunohistochemistry analysis was performed in midbrain sections from mice described in (D) to quantify 6-OHDA–induced dopaminergic neuronal loss in both injected (6-OHDA) and non-injected (control) sides (scale bar: 1 mm) (left panel). The total content of TH-positive somas was measured in midbrain sections covering the entire SNpc, in the non-injected (control) and injected (6-OHDA) side, for each group. Data represent the mean and standard error from 6-10 animals per group (right panel). (C) The expression of BiP was monitored in the brain obtained from AAV-UPRplus, AAV-ATF6f, AAV-XBP1s, or AAV-Mock injected animals for 6 months (upper panel) or 1 year (bottom panel) using immunohistochemistry with an anti-BiP antibody. (Scale bar: 100 μm). (H) Experimental design to evaluate the effects of UPRplus in an idiopathic PD model. Animals were injected with PFF α-synuclein into the striatum using brain stereotaxis. 1,5 months later, animals were injected with UPRplus or control vectors into the substantia nigra pars compacta (SNpc) followed by histological analysis 1,5 months later. (I) Transmission electronic microscopy of sonicated *α*-synuclein aggregated fibrils (Scale bar 100 nm). (J) Immunohistochemical analysis of the phosphorylated α-synuclein (p-α-syn) levels in SNpc region (scale bar: 100 μm) (left panel). Quantification of the p-α-syn levels (integrated density) covering SNpc region. Values are expressed as the average and standard error. Mock n = 9, UPRplus n = 9, XBP1s = 9, ATF6f n = 6 (right panel). Statistical analysis was performed using Dunnett’s multiple comparisons test (*: *p* < 0.05; **: *p* < 0.01; ***: *p* < 0.001).

We evaluated the impact of UPRplus in the motor coordination of animals injected with 6-OHDA using the cylinder test to evaluate locomotor asymmetry of injected mice. Administration of UPRplus in the SNpc reduced motor impairment caused by the 6-OHDA lesion (Figure 6D), observing a ~50% recovery (Figure S7A). These beneficial effects were not observed when equal titers of AAV-XBP1s or AAV-ATF6f were injected under the same experimental conditions (Figure 6D). As a control for the injection of 6-OHDA, the levels of striatal denervation were monitored using tyrosine hydroxylase (TH) staining. Similar levels of striatal denervation were detected in all experimental groups (near 80% loss) (Figure 6E), indicating the equivalent efficiency of the neurotoxin in all experimental groups.

To determine the possible neuroprotective effects of UPRplus *in vivo,* we quantified the number of dopaminergic neurons located in the entire SNpc region using serial sections images. A global reduction in the number of neurons was observed when UPRplus was administrated to the SNpc (Figures S7B and S7C). The administration of 6-OHDA led to a 40% reduction in the total number of dopaminergic neurons (Figure 6F). This neuronal loss was ameliorated in near 20% when AAV-UPRplus was delivered into the SNpc (Figures 6F, S7B and S7C). In contrast, the expression of XBP1s or ATF6f resulted in a slight protection when compared to UPRplus. Overall, no toxicity of the AAV-UPRplus construct was observed when injected into the SNpc (Figures S7 and S8).

To complement our results, we evaluated the possible effects of UPRplus on the levels of α-synuclein aggregation using an idiopathic model of PD that do not rely on overexpression (Figure 6H). This animal model is based on the aggregation and spreading of endogenous α-synuclein triggered by the intracerebral injection of preformed fibrils of recombinant α-synuclein (PFF)^96^. We confirmed the required size distribution of the fibrils by electron microscopy (Figure 6I) followed by quantification (Figure S9A), in addition to assess its aggregation capacity after exposure of primary cultures of cortical neurons to PFF (Figure S9B). Then, we injected α-synuclein PFF into the striatum of mice followed by the injection of AAVs 6 weeks later at the SNpc. This strategy resulted in the spreading of α-synuclein from the striatum to the SNpc as measured by increased levels of phosphorylated α-synuclein at Ser129 (a marker of its aggregation) (Figure 6J). Remarkably, the expression of UPRplus at the SNpc significantly reduced the content of phosphorylated α-synuclein. Under the same experimental conditions, ATF6f expression had a similar capacity to reduce α-synuclein aggregation, whereas XBP1s alone did not have a significant effect (Figure 6J).

To increase the translational value of this study, we also performed long-term experiments to determine the stability of the UPRplus construct and possible toxicity by analyzing animals after 6 or 12 months of injection. We observed low levels of toxicity and a detectable expression of UPRplus in all months tested (Figure S8A). Also, we were able to detect the presence of the UPRplus, XBP1s or ATF6f construct in the nucleus and the upregulation of BiP in the injected side (Figures 6G and S8B). After 1 year of injection, UPRplus presented a higher capacity to induce BiP at the SNpc when compared to XBP1s or ATF6f alone (Figures 6G).

Overall, these results suggest that our gene transfer strategy to deliver UPRplus into selective brain areas has a higher potency in reducing signs of neurodegeneration when compared to XBP1 s or ATF6f alone.

## Discussion

The UPR is the main cellular pathway governing adaptive mechanisms to reestablish proteostasis of the secretory pathway following an ER insult. To study the functional significance of the ATF6f/XBP1s heterodimer, we generated a strategy to enforce the expression of an artificial fusion construct that might increase the physical intramolecular interaction between both UPR transcription factors domains. Our results provide the first *in vivo* evidence indicating that the concerted action of XBP1s and ATF6s triggers transcriptional changes that are potentiated to enhance the capacity of cells to reduce abnormal protein aggregation and sustain cell survival in disease settings. Although both ATF6f and XBP1s control the upregulation of partially overlapping sets of target genes (i.e. ERAD components^13, 20^), the expression of UPRplus was highly selective for genes involved in protein folding and protein degradation.

Although the contribution of XBP1s and ATF6f to proteostasis balance is well established, organisms and tissues that are genetically ablated for these UPR transcription factors have divergent phenotypes^97^. Mice lacking XBP1 are embryonic lethal, and conditional deficiency of XBP1 have demonstrated central roles in plasma cell differentiation, the function of salivary glands, and the exocrine pathway, in addition to liver biology among other functions^14^. In contrast, mice lacking ATF6α develop normally^20^, do not show altered B cell function^98^, but the removal of the two mammalian homologs ATF6α and ATF6β is embryonically lethal^22, 99^. However, under experimental ER stress, ATF6α knockout animals develop liver steatosis resulting in lethality^100, 101^. These findings suggest that the combined function of XBP1s and ATF6f activities may have non-overlapping consequences on the proteostasis network in a tissue-specific manner.

Although UPR signaling has a dual role for the global ER stress response (induction of repair programs and apoptosis), gene expression reprogramming by XBP1s or ATF6f in mammals is exclusively linked to pro-survival outputs. Chronic ER stress has been linked to a series of degenerative conditions affecting the brain, in addition to metabolic diseases (obesity and diabetes), ischemia-reperfusion, eye disease, chronic inflammatory, among other pathological conditions^4, 7, 102^. Thus, the UPR, and more specifically ATF6f and XBP1s, represent interesting candidates for drug discovery, however most compounds available targeting these pathways are inhibitors^31^. In general, these small molecules have been developed to ablate the pro-survival effects of the UPR signaling in models of cancer because the pathway confers a selective force to drive oncogenic transformation and sustain tumor growth^103, 104^. Small molecules to activate and inhibit the PERK/eIF2α pathway have been also developed, however due to the complex nature of the effector outputs of this signaling branch (i.e. prosurvival, apoptosis, metabolism, global protein synthesis control, among others), side effects of such treatments are difficult to predict^105^. Small molecules that inhibit protein disulfide isomerases have been shown to activate ATF6, resulting in promising therapeutic effects in models of brain and heart ischemia-reperfusion^55, 83, 106^. XBP1s activators were recently reported, however these drugs were not tested in vivo^107^. Thus, gene therapy to deliver active UPR components into specific tissues is emerging as an attractive strategy to enforce UPR adaptive response and also target specific tissues in a restricted manner, avoiding the chronic and systemic administration of small molecules^37^.

Our results suggest that the combination of XBP1s and ATF6 as fusion proteins has broad potential in reducing abnormal protein aggregation. We choose HD and PD models for proof-of-concept because mHtt and α-synuclein expression perturb the function of the secretory pathway, and ER stress feedback to enhance abnormal protein aggregation^108^ (reviewed in^34, 82, 109^). Unbiased studies using yeast screenings demonstrated that α-synuclein abnormally interacts with Rab1, blocking ER to Golgi trafficking with resultant ER stress^110–112^. α-synuclein aggregates are also present at the ER lumen and form abnormal complexes with BiP^113–115^. Interactome studies revealed that mHtt block ERAD through a physical interaction, leading to chronic ER stress^116, 117^. Finally, gene expression profiling also demonstrated that ER stress is the major pathological signature triggered by 6-OHDA and other neurotoxins^91, 92^. UPRplus was effective in improving the survival of dopaminergic neurons at the SNpc in a pharmacological model of PD and reduced α-synuclein aggregation on an idiopathic PD model. In the context of gene therapy applications, PD is one of the main neurodegenerative diseases that promise positive outcomes for disease intervention because the neuronal populations affected are in part restricted to the SNpc, which is suitable for efficient transduction with recombinant AAVs^118, 119^. Multiple studies have demonstrated that ER stress is a major driver of dopaminergic neuron degeneration in PD^120, 121^. Markers of ER stress are detected in human postmortem tissues from PD patients^122, 123^, which can be even observed in incidental cases that presented Lewy body pathology^124^. Previous studies indicated that the ectopic expression of XBP1 into the SNpc using AAVs or lentiviruses provides partial protection against neurodegeneration in PD models^43, 44^. In our study XBP1s overexpression did not show significant protection in our PD model, which might be related to the AAV titer used, the time point analyzed and/or the use of different constructs (human cDNA, HA tag, no GFP marker, different purification and quantification systems). Our experiments were designed to see protection with UPRplus, and under the same conditions compare with the effects of expressing either XBP1s or ATF6f alone. Our aim was to develop a novel “tool reagent” that forces the heterodimerization between XBP1s and ATF6f. It is most likely that when the co-expression of XBP1 and ATF6f is performed, most of the protein complexes formed in the cell are homodimers between XBP1s or ATF6f, and a small fraction may form heterodimers. The affinities between homodimers are predicted to be higher than heterodimers, although this has not been tested. Our experimental approach is the first direct attempting to determine the biological activity of the heterodimer and also showed that it has a therapeutic potential. Previous data is correlative in terms of assigning a function to the ATF6-XBP1 s heterodimer. Our results indicate that UPRplus has a stronger effect in reducing neuronal loss in PD models, in addition to full-length mHtt aggregation when compared with XBP1s or ATF6f alone under the same experimental conditions. BiP is a major ER chaperone that globally enhances the capacity of cells to cope with ER stress^125^. The function of BiP has been proposed as a protective factor in PD models, and gene therapy to overexpress BiP using AAVs alleviated dopaminergic neuron loss and reduced α-synuclein aggregation *in vivo*^95, 126^. In addition, the delivery of AAV-BiP into the retina improved cell survival and functionality on an animal model of retinitis pigmentosa due to the expression of mutant rhodopsin^127^. Since BiP was one of the major genes induced by UPRplus identified in our unbiased proteomic screening, the ATF6f-XBP1s fusion protein might have major effects in promoting protein folding to sustain proteostasis. Overall, here we have designed and validated a novel and powerful tool to fine-tune gene expression and improve ER proteostasis with a therapeutic gain by exploiting the cooperation between two major UPR signaling branches.

## Materials and Methods

### Plasmid constructs and transfection

DNA sequences encode for the human ATF6f, XBP1s, and UPRplus were synthesized *de novo* and cloned with the HA epitope into the pAAV-CMV vector by Genewiz. The linker sequence corresponds to:

#### LFG

5’-CTA GGT GGT GGT GGT TCG GGT GGT GGT GGT TCG GGT GGT GGT GGT TCG GCG GCG GCG-3’

#### LAHA

5’-CTA GCG GAA GCG GCG GCG AAA GAA GCG GCG GCG AAA GAA GCG GCG GCG AAA GAA GCG GCG GCG AAA GCG GCG GCG-3’

#### LF

5’-CTA TTT AAT AAA GAA CAA CAA AAT GCG TTT TAT GAA ATA CTA CAT CTA CCG AAT CTA AAT GAA GAA CAA CGT AAT GGT TTT ATA CAA TCG CTA AAA GAT GAT CCG TCG CAA TCG GCG AAT CTA CTA GCG GAA GCG AAA AAA CTA AAT GAT GCG CAA GCG GCG GCG-3’.

pAAV-mHttQ85-mRFP contains the first 588 amino acids of the Htt gene with a tract of 85 glutamines, fused to mRFP were previously reported^38^. All transfections were performed using Effectene reagent (Qiagen) according to the manufacturer’s instructions. DNA was purified with Qiagen kits. Polyglutamine 79 track were in-frame N-terminal fusion of GFP previously described^128^. α-synuclein-WT-RFP vectors were provided by Dr. Hiroyoshi Ariga. siRNA pool for HSP90B1, HYOU1, PDIA4, HSPA5, PSBM7, and scramble (SCR) were purchased from Santa Cruz and transfections were made with Lipofectamine® RNAiMAX transfection Reagent from Invitrogen.

### Homology modeling of the heterodimer

PDB id 5T01, human c-Jun DNA binding domain homodimer in complex with methylated DNA, was selected as a template after an initial search with NCBI BLAST^129^ against all PDB^130^ protein sequences with both sequences. Global alignments were performed with NEEDLE^131^ using by-default parameters between each of the transcription factors and each of the template sequences obtained from PDBFINDER^132^. These alignments were hand-curated and then were employed as input for SCWRL V4^133^. For this analysis, we used the fragment 306-367 for the human ATF6f sequence (UniProtKB - P18850) and the fragment 70-131 for the human XBP1s sequence (UniProtKB - P17861).

### ESpritz based analysis of mobility

ESpritz is a predictor of disordered regions in protein sequences, one of its three versions were trained to predict amino acids that lack coordinates for all, are at least some, their atoms in X-ray solved protein structures. ESpritz employs BRNNs (Bidirectional Recurrent Neural Networks), a type of neural network that “reads” whole sequences to predict each of the elements in it, thus it considers the whole sequence to predict a property for each amino acid in it. ESpritz was run with by-default parameters using its web server. We analyzed the whole sequence of six UPRplus versions including the HA tag.

### Promoter region analysis

The promoter sequence was extracted 10kb before the transcription initiation site for the genes encoding for PDIA4, HSP90B1, HSPA5, PSMB7, and HYOU1. Transcription factor binding motifs (TFBMs) were searched for the human XBP1 transcription factors and human ATF6 using the CISBP library^134^ that contains information about transcription factors and their DNA binding domains. The motif format of this library was converted to MEME format, a format accepted by the FIMO program (‘Find Individual Motif Occurrences), which scans a set of sequences to search for individual matches to the motives provided.

### RNA isolation, RT-PCR and real-time PCR

Total RNA was prepared from tissues or cells placed in cold PBS using Trizol following the manufacturer’s instructions (Life Technologies). The cDNA was synthesized with SuperScript III reverse transcriptase (Life Technologies) using random primers p(dN)6 (Roche). Quantitative PCR reactions were performed in a BioRad CFX96 system employing the SYBRgreen fluorescent reagent (Applied Biosystems, USA). The relative amounts of mRNAs were calculated from the values of the comparative threshold cycle by using actin as a control. RT-PCR were performed using the following primers: For human: **HSPA5** 5’-GCCTGTATTTCTAGACCTGCC-3’ and 5’-TTCATCTTGCCAGCCAGTTG-3’, **CRELD2** 5’-ACTGAAGAAGGAGCACCCCAAC-3’ and 5’-CACACTCATCCACATCCACACA-3’, **SULF1** 5’-ATTCAAGGAGGCTGCTCAGG-3’ and 5’-TGTCATGCGTGAAGCAAGTG, **PDIA4** 5’-TGCCGCTAACAACCTGAGAG-3’ and 5’-TCCATGGCGAACTTCTTCCC-3’ **HSP90B1** TCCATATTCGTCAAACAGACCAC-3’ and 5’-CTGGGACTGGGAACTTATGAATG-3’, **PSMB7** 5’-TTTCTCCGCCCATACACAGTG-3’ and 5’-AGCACCTCAATCTCCAGAGGA-3’, **ANP32C** 5’-AACGACTACGGAGAAAACGTG-3’ and 5’-CCTTGTGGTCCCAGTAACAGC-3’, **beta Actin** 5’-GCGAGAAGATGACCCAGATC-3’ and 5’-CCAGTGGTACGGCCAGAGG-3’, For mouse: **HspA5** TCATCGGACGCACTTGGAA-3’ and 5’-CAACCACCTTGAATGGCAAGA-3’ **HerpUD1** 5’-CCCATACGTTGTGTAGCCAGA-3’ and 5’-GATGGTTTACGGCAAAGAGAAGT-3’, **Creld2** 5’-CAACACGGCCAGGAAGAATTT-3’ and 5’-CATGATCTCCAGAAGCCGGAT-3’ **Hyou1** 5’-TGCGCTTCCAGATCAGTCC-3’ and 5’-GGAGTAGTTCAGAACCATGCC-3’ **beta Actin** 5’-TACCACCATGTACCCAGGCA-3’ and 5’-CTCAGGAGGAG AATGATCTTGAT-3’, **EGFP** 5’-TCCTGGACGTAGCCTTCG-3’ and 5’-GGCAAGCTGACCCTGAAAGT-3’, **HA** 5’-TAGACGTAATCTGGAACATCG-3’.

### Microscopy, western blot and, filter trap analysis

Neuro2A and HEK293T cells were obtained from ATCC and maintained in Dulbecco’s modified Eagles medium supplemented with 5% fetal bovine serum. 3 × 105 cells were seeded in a 6-well plate and maintained by indicated times in DMEM cell culture media supplemented with 5% bovine fetal serum and non-essential amino acids. We visualized and quantified the formation of intracellular polyQ_79_-EGFP inclusions in living cells after transient transfection using epifluorescent microscopy. Intracellular inclusions were quantified using automatized macros done in Image J software. This Macros identify the cell total number using a intensity low threshold. The polyQ79 intracellular inclusions represent a saturated spot and were identified in the same Macros using a higher threshold. Protein aggregation was evaluated by western blot in total cell extracts prepared in 1% Triton X-100 in PBS containing proteases and phosphatases inhibitors (Roche). Protein quantification was performed with the Pierce BCA Protein Assay Kit (Thermo Scientific).

For western blot analysis, cells were collected and homogenized in RIPA buffer (20 mM Tris pH 8.0, 150 mM NaCl, 0.1% SDS, 0.5% Triton X-100) containing protease and phosphatase inhibitors (Roche). After sonication, protein concentration was determined in all experiments by micro-BCA assay (Pierce), and 25-100 μg of total protein was loaded onto 8 to 15 % SDS-PAGE mini gels (Bio-Rad Laboratories, Hercules, CA) prior transfer onto PVDF membranes. Membranes were blocked using PBS, 0.1% Tween-20 (PBST) containing 5% milk for 60 min at room temperature and then probed overnight with primary antibodies in PBS, 0.02% Tween-20 (PBST) containing 5% skimmed milk. The following primary antibodies and dilutions were used: anti-GFP 1:1,000 (Santa Cruz, Cat. n° SC-9996), anti-alpha-synuclein 1:1,000 (BD, Cat. n° 610787), anti-polyQ 1:1,000 (Sigma, Cat. n° P1874), anti-HSP90 1:2,000 (Santa Cruz, Cat. n° SC-13119), anti-GAPDH 1:2,000 (Santa Cruz, Cat. n° SC-365062) and anti-HA 1:500 (Santa Cruz, Cat. n°SC-805). Bound antibodies were detected with peroxidase-coupled secondary antibodies incubated for 2 h at room temperature and the ECL system.

For filter trap assays, protein extracts were diluted into a final concentration of SDS 1% and were subjected to vacuum filtration through a 96-well dot blot apparatus (Bio-Rad Laboratories, Hercules, USA) containing a 0.2 μM cellulose acetate membrane (Whatman, GE Healthcare) as described in (Torres et al., 2015). Membranes were then blocked using PBS, 0.1% Tween-20 (PBST) containing 5% milk and incubated with primary antibody at 4°C overnight. Image quantification was done with the Image Lab software from BioRad.

HEK293 DAX cells were maintained in Dulbecco’s modified Eagles medium supplemented with 5% fetal bovine serum. 3 × 105 cells were seeded in a 6-well plate and maintained by indicated times in DMEM cell culture media supplemented with 5% bovine fetal serum and non-essential amino acids. HEK293 DAX cells were treated by 16 hours with doxycycline (DOX) (1 uM) or Trimethoprim (TMP) (10 uM) to induce the XBP1s or ATF6f respectively.

### Electrophoretic mobility shift assay (EMSA)

The nuclear extract was performed using the NE-PER kit (Thermo Fisher Scientific). Electrophoretic mobility shift assay was performed using nuclear extracts obtained from HEK cells transiently transfected with pAAV-ATF6f-HA, pAAV-UPRplus-HA, or pAAV-empty (control). After 48 h, 10 μg of nuclear extracts were incubated with 200 fmol of 5’-biotin-labeled UPRE probe, 2 μl of 10 × binding buffer, 1 μl of poly dI/dC and 1 μL of 50% glycerol in a volume of 20 μl. For the competition assay, unlabeled or mutated probes were added to the reaction mixture 10 min before adding the labeled UPRE probe. The DNA-protein complexes were separated in a 6% non-denaturing polyacrylamide gel and were detected by western blot using an anti-biotin antibody.

### Quantitative proteomics

HEK cells in 6-well plates were infected with AAV-UPRplus, AAV-XBP1s, AAV-ATF6f, or AAV-empty for 48 h. Lysates were prepared in RIPA buffer (150 mM NaCl, 50 mM Tris pH 7.5, 1% Triton X-100, 0.5% sodium deoxycholate, and 0.1% SDS) containing proteases and phosphatases inhibitors cocktail (Roche). After extract sonication, protein concentration was determined by BCA (Thermo Fisher). For each sample, 20 μg of lysate was washed by chloroform/methanol precipitation. Samples for mass spectrometry analysis were prepared as described ^83^. Air-dried pellets were resuspended in 1% RapiGest SF (Waters) and brought up in 100 mM HEPES (pH 8.0). Proteins were reduced with 5mM Tris(2-carboxyethyl) phosphine hydrochloride (Thermo Fisher) for 30 min and alkylated with 10 mM iodoacetamide (Sigma Aldrich) for 30 min at ambient temperature and protected from light. Proteins were digested for 18 h at 37°C with 0.5 μg trypsin (Promega). After digestion, the peptides from each sample were reacted for 1 h with the appropriate TMT-NHS isobaric reagent (Thermo Fisher) in 40% (v/v) anhydrous acetonitrile and quenched with 0.4% ammonium bicarbonate for 1 h. Samples with different TMT labels were pooled and acidified with 5% formic acid. Acetonitrile was evaporated on a SpeedVac and debris was removed by centrifugation for 30 min at 18,000 × *g.* MuDPIT microcolumns were prepared as described ^135^. LCMS/MS analysis was performed using a Q Exactive mass spectrometer equipped with an EASY nLC 1000 (Thermo Fisher). MuDPIT experiments were performed by sequential injections of 0, 10, 20, 30, …, 100% buffer C (500 mM ammonium acetate in buffer A) and a final step of 90% buffer C / 10% buffer B (20% water, 80% acetonitrile, 0.1% formic acid, v/v/v) and each step followed by a gradient from buffer A (95% water, 5% acetonitrile, 0.1% formic acid) to buffer B. Electrospray was performed directly from the analytical column by applying a voltage of 2.5 kV with an inlet capillary temperature of 275°C. Data-dependent acquisition of MS/MS spectra was performed with the following settings: eluted peptides were scanned from 400 to 1800 m/z with a resolution of 30000 and the mass spectrometer in a data-dependent acquisition mode. The top ten peaks for each full scan were fragmented by HCD using a normalized collision energy of 30%, a 100 ms injection time, a resolution of 7500, and scanned from 100 to 1800 m/z. Dynamic exclusion parameters were 1 repeat count, 30 ms repeat duration. Peptide identification and protein quantification were performed using the Integrated Proteomics Pipeline Suite (IP2, Integrated Proteomics Applications, Inc., San Diego, CA) as described previously ^83^. TMT intensities were normalized to global peptide levels, summed for proteins and log2 transformed. Differences in protein expression were expressed as log2 fold changes between conditions and averaged across biological and technical replicates (Vehicle: n = 5, ATF6: n = 3, XBP1s: n = 3, UPRplus: n = 6). The vehicle and UPRplus samples contained 3 technical replicates that were distributed across 3 independent MuDPIT runs. Significance of expression changes and q values were evaluated in Graphpad Prism using multiple t-tests and false discovery rate (FDR) correction with two-stage step-up method of Benjamini, Krieger and Yekutieli. The mass spectrometry proteomics data have been deposited to the ProteomeXchange Consortium via the PRIDE^136^ partner repositor with the dataset identifier PXD022554.

### Adeno-associated viral vectors

All AAV (serotype 2) vectors were produced by triple transfection of 293 cells using a rep/cap plasmid and pHelper (Stratagene, La Jolla CA, USA), and purified by column affinity chromatography as previously described^38, 43, 46, 137^.

### Animals and surgical procedures

Adults male mice C57BL/6j (3-month-old) were injected with 2 μl of virus AAV-Htt588^Q95^-mRFP or were co-injected with 2 μl of each virus AAV-Htt588^Q95^-mRFP and AAV-XBP1s, AAV-ATF6f, AAV-UPRplus or AAV-empty (control) in the right striatum, using the following coordinates: +0.5 mm anterior, +2 mm lateral and −3 mm depth (according to the atlas of Franklin and Paxinos, Second Edition, 2001), with a 1 μl/min infusion rate. The titer virus used was 1 × 10^8^ viral genomes/ul (VGs) for each of them. After 2 weeks mice were euthanized, and brain tissues were dissected for western blot analysis.

We employed as HD model the full length mHtt transgenic mice with 128 CAG repetitions termed YAC128 ^87^ obtained from The Jackson Laboratory. Mice were injected bilaterally in the striatum with 1 μl per hemisphere of AAV-XBP1s, AAV-ATF6f, AAV-UPRplus or AAV-empty. The injection of AAVs suspension was performed at two points of the striatal region using a 5 μl Hamilton syringe (Hamilton) using the following coordinates: +0.7 mm anterior, +1.7 mm lateral and −3 to 3.2 mm depth, with a 1 μl/min infusion rate. Four weeks later, mice were euthanized, and brain tissues were dissected for western blot analysis.

For UPRplus overexpression in R6/2 mice, 1-month-old animals were used. For stereotactic injections, mice were anesthetized using isoflurane, and affixed to a mouse stereotaxic frame (David Kopf Instruments). Mice were injected bilaterally in the striatum with 1 μl per hemisphere of AAV-XBP1s, AAV-ATF6f, AAV-UPRplus or AAV-empty. The injection of AAVs suspension was performed at two points of the striatal region using a 5 μl Hamilton syringe (Hamilton) using the following coordinates: +0.7 mm anterior, +1.7 mm lateral and −3 to 3.2 mm depth, with a 1 μl/min infusion rate. After 6 weeks, mice were euthanized for histochemical analysis.

For UPRplus over-expression in the SNpc using AAVs, we used 3-month-old male C57BL/6 mice. We injected 2 μl of virus unilaterally in the right SNpc using the following coordinates: AP: −0,29 cm, ML: −0.13 cm, DV: −0.42 cm (according to the atlas of Franklin and Paxinos, Second Edition, 2001). The titer virus used was 1 × 10^8^ viral genomes/ul (VGs) for each of them. After 2 weeks of viral vectors injection, the injection of 6-OHDA was performed in a single point, injecting 8 μg in the right striatum using the following coordinates: AP: +0.07 cm, ML: −0.17 cm, DV: −0.31 cm (according to the atlas of Franklin and Paxinos Second Edition, 2001). Mice were euthanized 7 days after 6-OHDA injections for histological analysis.

For the generation of PD idiopathic model, recombinant mouse α-synuclein PFFs were generated as previously described^138^. Briefly, mouse α-Syn protein was dissolved in PBS, the pH adjusted to 7.5, subsequently the protein filtered through 100 kDa MW-cut-off filters and incubated with constant agitation (1000 rpm) for 5 days at 37°C. After incubation the pellet, containing the insoluble fibrils, was separated from the supernatant by ultracentrifugation (100’000 g, 30 min, 4°C), re-suspended in PBS, and the fibrils fragmented by sonication (5 sec, 20% amplitude, 1X pulse on and 1X pulse off, for four times on ice) to obtain smaller seeds, aliquoted and stored at −80°C until use. The presence of amyloid-like fibrils was characterized by transmission electron microscopy.

All experiments were performed in accordance with the guidelines set by the animal care and use committee of the Faculty of Medicine at the University of Chile, with approved animal experimentation protocol CBA#0488 FMUCH and CBA#0904 FMUCH.

### Motor test

The cylinder test was performed to evaluate spontaneous motor changes associated with dopamine depletion in the striatum of 6-OHDA injected mice. Animals were placed in a glass cylinder and the number of times the mouse touched the glass wall with each forepaw was recorded for 5 minutes using a video camera. An animal injected with 6-OHDA in the right striatum will touch more times with the opposite paw (contralateral) because the toxin induces striatal denervation of dopaminergic neurons and therefore dopamine depletion. The analysis was performed by a researcher blind for the experiment. The result is plotted as the percentage of contralateral touches relative to total touches with both forepaws.

### Tissue preparation and analysis

Mice were euthanized by CO_2_ narcosis, after which brains were rapidly removed and the ventral midbrain, containing the entire substantia nigra, striatum, and cortex from both hemispheres were promptly dissected on an ice-cold plastic dish. The tissue was homogenized in 100 μL of ice-cold 0.1 M PBS (pH 7.4) supplemented with a protease inhibitor cocktail (Roche). The homogenate was divided into two fractions for further total mRNA and protein extraction, followed by standard purification and quantification protocols. Protein extraction was performed in RIPA buffer (20 mM Tris pH 8.0, 150 mM NaCl, 0.1% SDS, 0.5% deoxycholate, and 0.5% Triton X-100) containing a protease and phosphatase inhibitor cocktail (Sigma-Aldrich). Sample quantification was performed with the Pierce BCA Protein Assay Kit (Thermo Scientific). For western blot analyses, samples were lysed, and extracts were loaded into SDS/PAGE gels and blotted onto PVDF membranes. Membranes were incubated with primary antibodies, followed by the incubation with secondary antibodies tagged with HRP. The following antibodies were used: Hsp90 (1:3,000; Santa Cruz Biotechnology), beta-actin (1:3,000; Santa Cruz Biotechnology), TH (1:2,000; Chemicon), HA (1:1,000; Abcam), GFP (1:3,000; Santa Cruz Biotechnology) and anti-α-synuclein antibody (1:1,000, BD Bioscience).

For RNA extraction and real-time PCR, total RNA was isolated from ventral midbrain (containing entire SNpc), striatum, and cortex. After cDNA production, real-time PCR was performed in a BioRad CFX96 system employing the SYBRgreen fluorescent reagent (Applied Biosystems, USA).

### Tissue Preparation and Histological Analysis

Mice were anesthetized and perfused through the ascending aorta with isotonic saline solution followed by ice-cold 4% paraformaldehyde in 0.1 M PBS (pH 7.4). Brains were frozen and coronal sections of 25 or 30 μm containing the rostral striatum and midbrain were cut on a cryostat (Leica, Germany). Free-floating midbrain and striatal tissue sections were stained following standard protocols.

For immunohistochemical analysis, sections were incubated overnight at 4°C in blocking solution with anti-TH (1:2,500; Chemicon), anti-phospho-α-synuclein 129 clone 81A antibody (1:1,000, Biolegend), anti-EM48 (1:500; Chemicon) or anti-HA (1:500; Roche) antibodies and developed with biotinylated secondary anti-rabbit (1:500; Vector Laboratories) and avidin-biotin-peroxidase complex (ABC Elite Kit; Vector Laboratories). For immunofluorescence analysis, sections were incubated overnight at 4°C in blocking solution with anti-TH (1:2,500; Chemicon) and anti-BiP (1:1,000; Calbiochem) antibodies and detected using secondary alexa-488 anti-rabbit and alexa-564 anti-mouse antibodies. Tissue staining was visualized with an inverted microscope Leica DMi8 for scanning complete sections.

Estimation of the number of TH-positive neurons stained by immunohistochemistry was performed manually by a researcher blind for the experiment. Results are expressed as the total number of TH-positive neurons per hemisphere. To determine the percentage of TH-positive cell loss in the SNpc of 6-OHDA injected mice, the number of dopaminergic cells in the injected and non-injected side was determined by counting in a blinded manner the total number of TH-positive cells in midbrain serial sections containing the entire SNpc (between the AP-0.29 and AP-0.35 cm coordinates)

Results are expressed as the percentage of TH-positive neurons in the injected side compared with the non-injected side. In addition, for striatum denervation quantifications, the images obtained by phase-contrast microscopy from serial sections covering the entire striatum were analyzed using ImageJ software (http://rsb.info.nih.gov/ij/). The total integrated density per hemisphere in the area was quantified. Results are expressed as the percentage of integrated density in the injected side compared with the non-injected side.

### Statistical analysis

Results were statistically compared using the one-way ANOVA for unpaired groups followed by Dunnett’s multiple comparisons test. A *p* value of < 0.05 was considered significant (*: *p* < 0.05; **: *p* < 0.01; ***: *p* < 0.001).

## Acknowledgements

This work was directly funded by FONDEF D11E1007 (CH/RLV). We also thank support from FONDECYT 1191003 (RLV), FONDAP program 15150012 (CH/RLV), Millennium Institute P09-015-F (CH/RLV), CONICYT-Brazil 441921/2016-7, FONDEF ID16I10223, and FONDECYT 1180186 (CH). In addition, we thank the support from the U.S. Air Force Office of Scientific Research FA9550-16-1-0384, US Office of Naval Research-Global (ONR-G) N62909-16-1-2003, (CH). We also thank FONDECYT 3150097 (PG-H), FONDECYT 1150069 (HGR), CONICYT Ph.D. fellowship 21160843 (PT-E), Programa de Apoyo a Centros con Financiamiento Basal AFB-170004 (SM), FONDECYT 1181089 (AM) and NIH NS092829, AG046495 (RLW).

## Author Contributions

Conceptualization, R.L.V., and C.H.; Methodology, R.L.V., and C.H.; Investigation, R.L.V., C.G., C.J., L.P., J.C., P.T-E., P.G-H., V.C., D.S., S.L., V.L., S.M., A.M. and C.R. Resources, R.L.V., R.L.W. and C.H.; Supervision, R.L.V., P.S., R.L.W. and C.H.; Writing – Original Draft, R.L.V., P.T-E., P.G-H., D.S. and C.H.; Writing – Review & Editing, P.T-E., R.L.V., D.S., P.G-H., R.L.W. and C.H. Project Administration, R.L.V. and C.H; Funding Acquisition, R.L.V. and C.H.

## Conflicts of interest

R.L.V. and C.H. protected the use of a gene therapy to deliver UPRplus into the brain to treat Parkinson’s and Huntington’s disease. UPRplus is a registered brand in Chile. The patent licensed to Handl Biotech, Belgium.

Title: **AAV/UPR-plus virus, UPR-plus fusion protein, genetic treatment method and use thereof in the treatment of neurodegenerative diseases, such as Parkinson’s disease and Huntington’s disease.**

Chilean application number: 3242-2015 (2015). Submitted date 4/11/2015.

International application number: PCT/CL2016/000070. Submitted date 4/11/2016.

International publication number: PCT WO2017075729A1. Publication date 11/05/2017.

PCT national phases: Australia, EPO and Japan.

Status: patent pending.

## Supplementary figure legends

**Table S1. Protein expression changes triggered by the expression of UPRplus, XBP1s, and ATF6f in HEK293T cells**. Quantitative proteomics was performed in protein extracts from HEK293T cells infected for 48 h with the following viral particles: AAV-UPRplus, AAV-XBP1s, AAV-ATF6f, or AAV-empty (vehicle). Fold change, q-value and Standard Deviation (SD) of 124 differential protein expression in each condition are showed. Color from red to blue indicates high to low expression.

**Table S2. Protein expression changes triggered by the expression of UPRplus, XBP1s, and ATF6f in HEK293T cells**. Quantitative proteomics was performed in protein extracts from HEK293T cells infected for 48 h with the following viral particles: AAV-UPRplus, AAV-XBP1s, AAV-ATF6f, or AAV-empty (vehicle). Raw data of differential protein expression in each condition are showed.

**Figure S1. Determination of ATF6f and XBP1s transcriptional activity and flexibility of UPRplus construct**. (A) The mRNA levels of ERdj4 (left panel) or GRP94 (right panel) genes were monitored in HEK293T cells infected with AAV-empty (control), AAV-UPRplus, AAV-XBP1s, AAV-ATF6f or AAV-XBP1s: AAV-ATF6f (1:1) viral particles. After 48 h the relative mRNA levels of indicated genes were measured by real-time PCR. All samples were normalized to β-*actin* levels. mRNA levels are expressed as fold increase over the value obtained in the control condition. (B) Quantification of the average disorder probability of primary protein structure considering the six fusion proteins used between ATF6f and XBP1s. The 0.1 default ESpritz threshold employed to annotate residues as disordered with a 5% False Positive Rate. (C) HEK-Rex DAX cells were treated by 16 hours with doxycycline (DOX) (1 uM), Trimethoprim (TMP) (10 uM) or both to induce the XBP1s or ATF6f respectively and transiently transfected with UPRplus, XBP1s, ATF6f or both XBP1s and ATF6f (A +X) vector. Relative mRNA levels of Sulf1 gene were measured by real-time PCR. All samples were normalized to β-*actin* levels. (D) polyQ_79_-EGFP detergent-insoluble aggregates were measured in cell extracts describe in C prepared in Triton X100 by western blot (right panel). Levels of Hsp90 were measured as the loading control. Left panel: high molecular weight (HMW) polyQ_79_-EGFP aggregates were quantified.

**Figure S2. Expression levels and in silico structural analysis of UPRplus.** (A) HEK293T cells were transiently transfected with HA-tagged expression vectors for UPRplus, ATF6f, XBP1s, or empty vector (control). After 48 h, cytosolic and nuclear extracts were analyzed by western blot using an anti-HA antibody. (B) The amino acid sequence of UPRplus including, ATF6f (blue), linker (yellow), and XBP1s (red) sequence. The putative dimerization domains are highlighted in a black box. Mutated residues used in figure 2 are underlined in green. (C) Alignment of the putative dimerization domain sequences for ATF6f (upper panel) or XBP1s (lower panel). Asterisks represent the conserved residues and mutated residues are highlighted in red. (D) The three-dimensional model of a heterodimer of XBP1s and ATF6f attached to a DNA motif (see methods). The ATF6f chain corresponds to 306-367 residues and XBP1s chain to 70-131 residues. The XBP1s chain appears in red and the ATF6f chain is shown in blue with the residues K122, K315, N316, and R317 highlighted. The position of the linker (yellow) and the rest of the sequence are indicated using lines. (E) HEK293T cells were transiently transfected with the UPRplus WT, UPRplus K122K, UPRplus K315T, UPRplus N316A, or UPRplus R3177 constructs and after 48 h were analyzed by western blot using an anti-HA antibody (upper panel). Bottom panel: Levels of Hsp90 were monitored as a loading control. (F) HEK293T cells were transiently transfected with the UPRplus WT, UPRplus K122K, UPRplus K315T, UPRplus N316A, or UPRplus R3177 constructs and after 48 h were analyzed by immunofluorescence using an anti-HA antibody (green). Co-staining with the nuclear marker Hoechst (blue) was performed. Scale bar 20 μm.

**Figure S3. The activity of UPRplus depends on the dimer interphase and DNA binding domain.** (A) Higher magnification images of Neuro2A cells transiently co-transfected with expression vectors for polyQ_79_-EGFP together with empty vector (control), UPRplus WT, or the UPRplus mutants K122L, K315T, N316A, or R317A. After 48 h of expression, the accumulation of polyQ_79_-EGFP intracellular inclusions was visualized by fluorescence microscopy. Scale bar, 20 μm. (B) Neuro2A cells were transiently co-transfected with expression vectors for polyQ_79_-EGFP together with empty vector (control), UPRplus WT, or the mutants K122L, N316A, R317A, or K315T. After 48 h, high molecular weight (HMW) polyQ_79_-EGFP species were measured in cell extracts prepared in Triton X100 using western blot with an anti-GFP antibody (upper panel). Bottom panel: Levels of Hsp90 were monitored as a loading control.

**Figure S4. Analysis of the promoter regions of genes regulated by UPRplus.** (A) The motif pattern analysis in the promoter region showing that the canonical ERSE I and ERSE II are present in the promoters of the five top genes upregulated by UPRplus. The promoter sequences spanning 15 kb upstream and 2 kb downstream of the transcriptional start site (TSS) were examined. The consensus sequence is shown in the indicated boxes. The blue box is the consensus sequence to which NF-Y binds and the red box is the consensus sequence to which ATF6 binds. (B) Frequency of transcription factor binding motifs (TFBMs) of transcription factors XBP1s and ATF6f at 10,000 bp upstream of the transcription start site. (C) Representation of the promoter region 10,000 bp upstream of the TSS for the five UPRplus upregulated genes in the positive and negative strands. Vertical marks correspond to the presence of specific motifs for XBP1s in red and ATF6f in blue. (D) Graphical representation of a nucleic acid multiple sequence alignment of motifs as sequence logo to ATF6f, XBP1 s, and both ATF6f/XBP1s, indicating the relative frequency of each nucleic acid in the motif.

**Figure S5. Validation of the knockdown experiments for UPRplus-regulated genes.** (A) HEK293T cells were transiently transfected with siRNA against indicated genes or scrambled siRNA (siSCR) as control. After 48 h, the expression levels of the indicated mRNAs were measured by real-time RT-PCR. All samples were normalized to β-actin levels. mRNA levels are expressed as fold increase over the value obtained in the control condition (siSCR). (B) After 48 h, the EGFP expression levels were measured by real-time PCR to control transfection efficiency. All samples were normalized to β-actin levels. mRNA levels are expressed as fold increase over the value obtained in the control condition (siSCR). In all experiments the mean and standard error is presented of three independent experiments. Statistical analysis was performed using Dunnett’s multiple comparisons test (**: *p* < 0.01; ***: *p* < 0.001).

**Figure S6. Validation of UPRplus expression in neurons.** (A) Neuro2A cells were transiently co-transfected with expression vectors for polyQ_79_-EGFP and XBP1s, ATF6f, UPRplus, or empty vector (Control). After 24 h, expression levels of human ATF6f and XBP1s were measured by real-time RT-PCR. All samples were normalized to β-actin levels. (B) Validation of UPRplus, XBP1s, or ATF6f expression in primary cortical neurons. Primary cortical neurons were infected at 1 day in vitro (DIV) with adeno-associated virus (AAV) encoding for UPRplus, XBP1s, ATF6f, or empty vector (control). After 6 DIV, expression levels of human ATF6f and XBP1s were measured by real-time RT-PCR. All samples were normalized to β-actin levels. (C) Validation of UPRplus, XBP1s, or ATF6f expression in YAC128 mice injected. Three-months old YAC128 mice were injected into the striatum with AAV-UPRplus, AAV-ATF6f, AAV-XBP1s, or AAV-Mock vector (control) using unilateral stereotaxis surgery. Four weeks later, the striatum was dissected and human XBP1s and ATF6f levels analyzed by PCR in each group.

**Figure S7. Effects of UPRplus expression on the survival of dopaminergic neurons after exposure to 6-OHDA.** (A) Analysis of the percentage of recovery in the cylinder test performance of animals treated with AAV-UPRplus. (B-C) Histograms show the number of TH-positive neurons of injected and non-injected sides in 25 μm midbrain serial sections separated by 100 μm and covering the entire SNpc. The number of serial sections indicates the orientation from anterior to posterior from animals injected with AAV-Mock (B) or AAV-UPRplus (C). Data represent the mean and standard error from 8 animals per group Statistical analysis was performed using Dunnett’s multiple comparisons test. *: *p* < 0.05, **: *p* < 0.01.

**Figure S8. Determination of safety and stability of UPRplus expression in the brain.** (A) The expression of TH was performed by immunohistochemistry in midbrain sections from mice injected with AAV-UPRplus, AAV-ATF6f, AAV-XBP1s, or AAV-Mock for 1 year. The total content of TH-positive somas was measured in midbrain sections covering the entire SNpc, in the non-injected (control) and injected side, for each group. Data represent the mean and standard error from 4 animals per group. (B) The expression of UPRplus was monitored in the brain obtained from AAV-UPRplus, AAV-XBP1s-HA, AAV-ATF6f-HA or AAV-Mock injected animals for 1 year using immunohistochemistry with an anti-HA antibody. (Scale bar: 1 mm, upper panel and 100 μm, bottom panel).

**Figure S9. Characterization of α-synuclein preformed fibrils.** (A) Frequency distribution of α-synuclein preformed fibrils (PFFs) length by transmission electron microscopy. Histogram of distribution for relative abundance of post-sonicated fibrils. 600 structures were counted. (B) Primary cortical neurons were treated with PBS (Control) (left panel) or with α-synuclein PFF (1 ng) at 7 day in vitro (DIV) (middle panel). After 7 DIV the phosphorylated form of α-synuclein (p-α-syn) (green), microtubule associate protein 2 (MAP2) (red) and nuclear staining with DAPI (blue) were detected by immunofluorescence. Higher magnification of middle panel (right panel). Scale bar: 50 μm (left and middle panel), 20 μm (right panel).

**Figure S10. Analysis of mHtt levels in a viral HD model.** (A-B) Three-month-old wild type mice were co-injected into the striatum by stereotaxis with a mixture of AAVs encoding a mHtt construct (Htt588^Q95^-mRFP) together with AAV-UPRplus, AAV-XBP1s, AAV-ATF6f, or AAV-Mock (control). Animals were then euthanized 2 weeks post-injection and brain striatum tissue was dissected for western blot analysis using an anti-polyQ antibody. β-actin levels were monitored as a loading control (bottom panel).

**Figure S11. Validation of the proteomic analysis.** The mRNA levels of selected UPR-upregulated genes were monitored in HEK293T cells infected with AAV-empty (control), AAV-XBP1s or AAV-ATF6f viral particles. After 48 h the relative mRNA levels of indicated genes were measured by real-time PCR. As a positive control, cells were treated with 1 μg/ml tunicamycin for 8 h (Tm). All samples were normalized to β-*actin* levels.

## References

1. Balch, WE, Morimoto, RI, Dillin, A, and Kelly, JW (2008). Adapting proteostasis for disease intervention. Science 319: 916–919.

2. Hiramatsu, N, Chiang, WC, Kurt, TD, Sigurdson, CJ, and Lin, JH (2015). Multiple Mechanisms of Unfolded Protein Response-Induced Cell Death. The American journal of pathology 185: 1800–1808.

3. Walter, P, and Ron, D (2011). The unfolded protein response: from stress pathway to homeostatic regulation. Science 334: 1081–1086.

4. Wang, M, and Kaufman, RJ (2016). Protein misfolding in the endoplasmic reticulum as a conduit to human disease. Nature 529: 326–335.

5. Hetz, C, Zhang, K, and Kaufman, RJ (2020). Mechanisms, regulation and functions of the unfolded protein response. Nat Rev Mol Cell Biol 21: 421–438.

6. Urra, H, Dufey, E, Lisbona, F, Rojas-Rivera, D, and Hetz, C (2013). When ER stress reaches a dead end. Biochim Biophys Acta.

7. Oakes, SA, and Papa, FR (2015). The role of endoplasmic reticulum stress in human pathology. Annu Rev Pathol 10: 173–194.

8. Karagoz, GE, Acosta-Alvear, D, and Walter, P (2019). The Unfolded Protein Response: Detecting and Responding to Fluctuations in the Protein-Folding Capacity of the Endoplasmic Reticulum. Cold Spring Harb Perspect Biol 11.

9. Yoshida, H, Matsui, T, Yamamoto, A, Okada, T, and Mori, K (2001). XBP1 mRNA is induced by ATF6 and spliced by IRE1 in response to ER stress to produce a highly active transcription factor. Cell 107: 881–891.

10. Lee, K, Tirasophon, W, Shen, X, Michalak, M, Prywes, R, Okada, T, et al. (2002). IRE1-mediated unconventional mRNA splicing and S2P-mediated ATF6 cleavage merge to regulate XBP1 in signaling the unfolded protein response. Genes Dev 16: 452–466.

11. Calfon, M, Zeng, H, Urano, F, Till, JH, Hubbard, SR, Harding, HP, et al. (2002). IRE1 couples endoplasmic reticulum load to secretory capacity by processing the XBP-1 mRNA. Nature 415: 92–96.

12. Acosta-Alvear, D, Zhou, Y, Blais, A, Tsikitis, M, Lents, NH, Arias, C, et al. (2007). XBP1 controls diverse cell type- and condition-specific transcriptional regulatory networks. Mol Cell 27: 53–66.

13. Lee, AH, Iwakoshi, NN, and Glimcher, LH (2003). XBP-1 regulates a subset of endoplasmic reticulum resident chaperone genes in the unfolded protein response. Molecular and cellular biology 23: 7448–7459.

14. Hetz, C, Martinon, F, Rodriguez, D, and Glimcher, LH (2011). The unfolded protein response: integrating stress signals through the stress sensor IRE1alpha. Physiol Rev 91: 1219–1243.

15. Hollien, J, and Weissman, JS (2006). Decay of endoplasmic reticulum-localized mRNAs during the unfolded protein response. Science 313: 104–107.

16. Hollien, J, Lin, JH, Li, H, Stevens, N, Walter, P, and Weissman, JS (2009). Regulated Ire1-dependent decay of messenger RNAs in mammalian cells. J Cell Biol 186: 323–331.

17. Han, D, Lerner, AG, Vande Walle, L, Upton, JP, Xu, W, Hagen, A, et al. (2009). IRE1alpha kinase activation modes control alternate endoribonuclease outputs to determine divergent cell fates. Cell 138: 562–575.

18. Urra, H, Pihan, P, and Hetz, C (2020). The UPRosome - decoding novel biological outputs of IRE1alpha function. Journal of cell science 133.

19. Asada, R, Kanemoto, S, Kondo, S, Saito, A, and Imaizumi, K (2011). The signalling from endoplasmic reticulum-resident bZIP transcription factors involved in diverse cellular physiology. Journal of biochemistry 149: 507–518.

20. Yamamoto, K, Sato, T, Matsui, T, Sato, M, Okada, T, Yoshida, H, et al. (2007). Transcriptional induction of mammalian ER quality control proteins is mediated by single or combined action of ATF6alpha and XBP1. Developmental cell 13: 365–376.

21. Haze, K, Yoshida, H, Yanagi, H, Yura, T, and Mori, K (1999). Mammalian transcription factor ATF6 is synthesized as a transmembrane protein and activated by proteolysis in response to endoplasmic reticulum stress. Molecular biology of the cell 10: 3787–3799.

22. Yamamoto, K, Sato, T, Matsui, T, Sato, M, Okada, T, Yoshida, H, et al. (2007). Transcriptional induction of mammalian ER quality control proteins is mediated by single or combined action of ATF6alpha and XBP1. Developmental cell 13: 365–376.

23. Hillary, RF, and FitzGerald, U (2018). A lifetime of stress: ATF6 in development and homeostasis. J Biomed Sci 25: 48.

24. Sudhakar, A, Ramachandran, A, Ghosh, S, Hasnain, SE, Kaufman, RJ, and Ramaiah, KV (2000). Phosphorylation of serine 51 in initiation factor 2 alpha (eIF2 alpha) promotes complex formation between eIF2 alpha(P) and eIF2B and causes inhibition in the guanine nucleotide exchange activity of eIF2B. Biochemistry 39: 12929–12938.

25. Harding, HP, Zhang, Y, Bertolotti, A, Zeng, H, and Ron, D (2000). Perk is essential for translational regulation and cell survival during the unfolded protein response. Mol Cell 5: 897–904.

26. Harding, HP, Zhang, Y, and Ron, D (1999). Protein translation and folding are coupled by an endoplasmic-reticulum-resident kinase. Nature 397: 271–274.

27. Walter, P, and Ron, D (2011). The unfolded protein response: from stress pathway to homeostatic regulation. Science 334: 1081–1086.

28. Han, J, Back, SH, Hur, J, Lin, YH, Gildersleeve, R, Shan, J, et al. (2013). ER-stress-induced transcriptional regulation increases protein synthesis leading to cell death. Nat Cell Biol 15: 481–490.

29. Tabas, I, and Ron, D (2011). Integrating the mechanisms of apoptosis induced by endoplasmic reticulum stress. Nat Cell Biol 13: 184–190.

30. Bettigole, SE, and Glimcher, LH (2015). Endoplasmic reticulum stress in immunity. Annu Rev Immunol 33: 107–138.

31. Hetz, C, Axten, JM, and Patterson, JB (2019). Pharmacological targeting of the unfolded protein response for disease intervention. Nature chemical biology 15: 764–775.

32. Soto, C, and Pritzkow, S (2018). Protein misfolding, aggregation, and conformational strains in neurodegenerative diseases. Nat Neurosci 21: 1332–1340.

33. Scheper, W, and Hoozemans, JJ (2015). The unfolded protein response in neurodegenerative diseases: a neuropathological perspective. Acta neuropathologica 130: 315–331.

34. Hetz, C, and Saxena, S (2017). ER stress and the unfolded protein response in neurodegeneration. Nature reviews Neurology 13: 477–491.

35. Smith, HL, and Mallucci, GR (2016). The unfolded protein response: mechanisms and therapy of neurodegeneration. Brain: a journal of neurology.

36. Morris, G, Puri, BK, Walder, K, Berk, M, Stubbs, B, Maes, M, et al. (2018). The Endoplasmic Reticulum Stress Response in Neuroprogressive Diseases: Emerging Pathophysiological Role and Translational Implications. Molecular neurobiology 55: 8765–8787.

37. Valenzuela, V, Jackson, KL, Sardi, SP, and Hetz, C (2018). Gene Therapy Strategies to Restore ER Proteostasis in Disease. Molecular therapy: the journal of the American Society of Gene Therapy 26: 1404–1413.

38. Zuleta, A, Vidal, RL, Armentano, D, Parsons, G, and Hetz, C (2012). AAV-mediated delivery of the transcription factor XBP1s into the striatum reduces mutant Huntingtin aggregation in a mouse model of Huntington’s disease. Biochem Biophys Res Commun 420: 558–563.

39. Casas-Tinto, S, Zhang, Y, Sanchez-Garcia, J, Gomez-Velazquez, M, Rincon-Limas, DE, and Fernandez-Funez, P (2011). The ER stress factor XBP1s prevents amyloid-beta neurotoxicity. Hum Mol Genet 20: 2144–2160.

40. Loewen, CA, and Feany, MB (2010). The unfolded protein response protects from tau neurotoxicity in vivo. PLoS One 5.

41. Waldherr, SM, Strovas, TJ, Vadset, TA, Liachko, NF, and Kraemer, BC (2019). Constitutive XBP-1s-mediated activation of the endoplasmic reticulum unfolded protein response protects against pathological tau. Nat Commun 10: 4443.

42. Cisse, M, Duplan, E, Lorivel, T, Dunys, J, Bauer, C, Meckler, X, et al. (2017). The transcription factor XBP1s restores hippocampal synaptic plasticity and memory by control of the Kalirin-7 pathway in Alzheimer model. Mol Psychiatry 22: 1562–1575.

43. Valdes, P, Mercado, G, Vidal, RL, Molina, C, Parsons, G, Court, FA, et al. (2014). Control of dopaminergic neuron survival by the unfolded protein response transcription factor XBP1. Proc Natl Acad Sci U S A 111: 6804–6809.

44. Sado, M, Yamasaki, Y, Iwanaga, T, Onaka, Y, Ibuki, T, Nishihara, S, et al. (2009). Protective effect against Parkinson’s disease-related insults through the activation of XBP1. Brain Res 1257: 16–24.

45. Hu, Y, Park, KK, Yang, L, Wei, X, Yang, Q, Cho, KS, et al. (2012). Differential effects of unfolded protein response pathways on axon injury-induced death of retinal ganglion cells. Neuron 73: 445–452.

46. Valenzuela, V, Collyer, E, Armentano, D, Parsons, GB, Court, FA, and Hetz, C (2012). Activation of the unfolded protein response enhances motor recovery after spinal cord injury. Cell Death Dis 3: e272.

47. Oñate, M, Catenaccio, A, Martínez, G, Armentano, D, Parsons, G, Kerr, B, et al. (2016). Activation of the unfolded protein response promotes axonal regeneration after peripheral nerve injury. Scientific reports 6: 21709–21709.

48. Schiattarella, GG, Altamirano, F, Tong, D, French, KM, Villalobos, E, Kim, SY, et al. (2019). Nitrosative stress drives heart failure with preserved ejection fraction. Nature 568: 351–356.

49. Wang, ZV, Deng, Y, Gao, N, Pedrozo, Z, Li, DL, Morales, CR, et al. (2014). Spliced X-box binding protein 1 couples the unfolded protein response to hexosamine biosynthetic pathway. Cell 156: 1179–1192.

50. Deng, Y, Wang, ZV, Tao, C, Gao, N, Holland, WL, Ferdous, A, et al. (2013). The Xbp1s/GalE axis links ER stress to postprandial hepatic metabolism. J Clin Invest 123: 455-468.

51. Zhou, Y, Lee, J, Reno, CM, Sun, C, Park, SW, Chung, J, et al. (2011). Regulation of glucose homeostasis through a XBP-1-FoxO1 interaction. Nat Med 17: 356–365.

52. Naranjo, JR, Zhang, H, Villar, D, Gonzalez, P, Dopazo, XM, Moron-Oset, J, et al. (2016). Activating transcription factor 6 derepression mediates neuroprotection in Huntington disease. J Clin Invest 126: 627–638.

53. Kazaz, IO, Demir, S, Yulug, E, Colak, F, Bodur, A, Yaman, SO, et al. (2019). N-acetylcysteine protects testicular tissue against ischemia/reperfusion injury via inhibiting endoplasmic reticulum stress and apoptosis. J Pediatr Urol 15: 253 e251-253 e258.

54. Glembotski, CC, Rosarda, JD, and Wiseman, RL (2019). Proteostasis and Beyond: ATF6 in Ischemic Disease. Trends Mol Med 25: 538–550.

55. Blackwood, EA, Azizi, K, Thuerauf, DJ, Paxman, RJ, Plate, L, Kelly, JW, et al. (2019). Pharmacologic ATF6 activation confers global protection in widespread disease models by reprograming cellular proteostasis. Nat Commun 10: 187.

56. Jin, JK, Blackwood, EA, Azizi, K, Thuerauf, DJ, Fahem, AG, Hofmann, C, et al. (2017). ATF6 Decreases Myocardial Ischemia/Reperfusion Damage and Links ER Stress and Oxidative Stress Signaling Pathways in the Heart. Circ Res 120: 862–875.

57. Yamamoto, K, Yoshida, H, Kokame, K, Kaufman, RJ, and Mori, K (2004). Differential contributions of ATF6 and XBP1 to the activation of endoplasmic reticulum stress-responsive cis-acting elements ERSE, UPRE and ERSE-II. Journal of biochemistry 136: 343–350.

58. Yoshida, H, Haze, K, Yanagi, H, Yura, T, and Mori, K (1998). Identification of the cis-acting endoplasmic reticulum stress response element responsible for transcriptional induction of mammalian glucose-regulated proteins. Involvement of basic leucine zipper transcription factors. J Biol Chem 273: 33741–33749.

59. Yoshida, H, Matsui, T, Hosokawa, N, Kaufman, RJ, Nagata, K, and Mori, K (2003). A time-dependent phase shift in the mammalian unfolded protein response. Developmental cell 4: 265–271.

60. Shoulders, MD, Ryno, LM, Genereux, JC, Moresco, JJ, Tu, PG, Wu, C, et al. (2013). Stress-independent activation of XBP1s and/or ATF6 reveals three functionally diverse ER proteostasis environments. Cell reports 3: 1279–1292.

61. Shaffer, AL, Shapiro-Shelef, M, Iwakoshi, NN, Lee, AH, Qian, SB, Zhao, H, et al. (2004). XBP1, downstream of Blimp-1, expands the secretory apparatus and other organelles, and increases protein synthesis in plasma cell differentiation. Immunity 21: 8193.

62. Bommiasamy, H, Back, SH, Fagone, P, Lee, K, Meshinchi, S, Vink, E, et al. (2009). ATF6alpha induces XBP1-independent expansion of the endoplasmic reticulum. Journal of cell science 122: 1626–1636.

63. Shoulders, MD, Ryno, LM, Genereux, JC, Moresco, JJ, Tu, PG, Wu, C, et al. (2013). Stress-Independent Activation of XBP1s and/or ATF6 Reveals Three Functionally Diverse ER Proteostasis Environments. Cell reports 3: 1279–1292.

64. Hetz, C (2012). The unfolded protein response: controlling cell fate decisions under ER stress and beyond. Nat Rev Mol Cell Biol 13: 89–102.

65. Ameri, K, and Harris, AL (2008). Activating transcription factor 4. Int J Biochem Cell Biol 40: 14–21.

66. Liu, J, Ibi, D, Taniguchi, K, Lee, J, Herrema, H, Akosman, B, et al. (2016). Inflammation Improves Glucose Homeostasis through IKKbeta-XBP1s Interaction. Cell 167: 1052–1066 e1018.

67. Chen, X, Iliopoulos, D, Zhang, Q, Tang, Q, Greenblatt, MB, Hatziapostolou, M, et al. (2014). XBP1 promotes triple-negative breast cancer by controlling the HIF1alpha pathway. Nature 508: 103–107.

68. Luo, R, Lu, JF, Hu, Q, and Maity, SN (2008). CBF/NF-Y controls endoplasmic reticulum stress induced transcription through recruitment of both ATF6(N) and TBP. Journal of cellular biochemistry 104: 1708–1723.

69. Gkogkas, C, Middleton, S, Kremer, AM, Wardrope, C, Hannah, M, Gillingwater, TH, et al. (2008). VAPB interacts with and modulates the activity of ATF6. Hum Mol Genet 17: 1517–1526.

70. Robinson, CR, and Sauer, RT (1998). Optimizing the stability of single-chain proteins by linker length and composition mutagenesis. Proc Natl Acad Sci U S A 95: 5929–5934.

71. Arai, R, Ueda, H, Kitayama, A, Kamiya, N, and Nagamune, T (2001). Design of the linkers which effectively separate domains of a bifunctional fusion protein. Protein engineering 14: 529–532.

72. Marqusee, S, and Baldwin, RL (1987). Helix stabilization by Glu-…Lys+ salt bridges in short peptides of de novo design. Proc Natl Acad Sci U S A 84: 8898–8902.

73. Walsh, I, Martin, AJ, Di Domenico, T, and Tosatto, SC (2012). ESpritz: accurate and fast prediction of protein disorder. Bioinformatics 28: 503–509.

74. Ono, SJ, Liou, HC, Davidon, R, Strominger, JL, and Glimcher, LH (1991). Human Xbox-binding protein 1 is required for the transcription of a subset of human class II major histocompatibility genes and forms a heterodimer with c-fos. Proc Natl Acad Sci U S A 88: 4309–4312.

75. Hai, TW, Liu, F, Coukos, WJ, and Green, MR (1989). Transcription factor ATF cDNA clones: an extensive family of leucine zipper proteins able to selectively form DNA-binding heterodimers. Genes Dev 3: 2083–2090.

76. Shuda, M, Kondoh, N, Imazeki, N, Tanaka, K, Okada, T, Mori, K, et al. (2003). Activation of the ATF6, XBP1 and grp78 genes in human hepatocellular carcinoma: a possible involvement of the ER stress pathway in hepatocarcinogenesis. Journal of hepatology 38: 605–614.

77. Acharya, A, Rishi, V, Moll, J, and Vinson, C (2006). Experimental identification of homodimerizing B-ZIP families in Homo sapiens. Journal of structural biology 155: 130–139.

78. Pogenberg, V, Consani Textor, L, Vanhille, L, Holton, SJ, Sieweke, MH, and Wilmanns, M (2014). Design of a bZip transcription factor with homo/heterodimer-induced DNA-binding preference. Structure 22: 466–477.

79. Seldeen, KL, McDonald, CB, Deegan, BJ, and Farooq, A (2009). Single nucleotide variants of the TGACTCA motif modulate energetics and orientation of binding of the Jun-Fos heterodimeric transcription factor. Biochemistry 48: 1975–1983.

80. Glover, JN, and Harrison, SC (1995). Crystal structure of the heterodimeric bZIP transcription factor c-Fos-c-Jun bound to DNA. Nature 373: 257–261.

81. Agre, P, Johnson, PF, and McKnight, SL (1989). Cognate DNA binding specificity retained after leucine zipper exchange between GCN4 and C/EBP. Science 246: 922–926.

82. Smith, HL, and Mallucci, GR (2016). The unfolded protein response: mechanisms and therapy of neurodegeneration. Brain 139: 2113–2121.

83. Plate, L, Cooley, CB, Chen, JJ, Paxman, RJ, Gallagher, CM, Madoux, F, et al. (2016). Small molecule proteostasis regulators that reprogram the ER to reduce extracellular protein aggregation. eLife 5.

84. Kokame, K, Kato, H, and Miyata, T (2001). Identification of ERSE-II, a new cis-acting element responsible for the ATF6-dependent mammalian unfolded protein response. J Biol Chem 276: 9199–9205.

85. Wang, S, Hu, B, Ding, Z, Dang, Y, Wu, J, Li, D, et al. (2018). ATF6 safeguards organelle homeostasis and cellular aging in human mesenchymal stem cells. Cell Discov 4: 2.

86. Garcia-Huerta, P, Troncoso-Escudero, P, Wu, D, Thiruvalluvan, A, Cisternas-Olmedo, M, Henriquez, DR, et al. (2020). Insulin-like growth factor 2 (IGF2) protects against Huntington’s disease through the extracellular disposal of protein aggregates. Acta neuropathologica.

87. Slow, EJ, van Raamsdonk, J, Rogers, D, Coleman, SH, Graham, RK, Deng, Y, et al. (2003). Selective striatal neuronal loss in a YAC128 mouse model of Huntington disease. Hum Mol Genet 12: 1555–1567.

88. Mangiarini, L, Sathasivam, K, Seller, M, Cozens, B, Harper, A, Hetherington, C, et al. (1996). Exon 1 of the HD gene with an expanded CAG repeat is sufficient to cause a progressive neurological phenotype in transgenic mice. Cell 87: 493–506.

89. Martinez, A, Lopez, N, Gonzalez, C, and Hetz, C (2019). Targeting of the unfolded protein response (UPR) as therapy for Parkinson’s disease. Biol Cell 111: 161–168.

90. Silva, RM, Ries, V, Oo, TF, Yarygina, O, Jackson-Lewis, V, Ryu, EJ, et al. (2005). CHOP/GADD153 is a mediator of apoptotic death in substantia nigra dopamine neurons in an in vivo neurotoxin model of parkinsonism. J Neurochem 95: 974–986.

91. Ryu, EJ, Harding, HP, Angelastro, JM, Vitolo, OV, Ron, D, and Greene, LA (2002). Endoplasmic reticulum stress and the unfolded protein response in cellular models of Parkinson’s disease. J Neurosci 22: 10690–10698.

92. Holtz, WA, and O’Malley, KL (2003). Parkinsonian mimetics induce aspects of unfolded protein response in death of dopaminergic neurons. J Biol Chem 278: 19367–19377.

93. Castillo, V, Mercado, G, and Hetz, C (2015). Gene therapy in Parkinson’s disease: targeting the endplasmic reticulum proteostasis network. Neural regeneration research 10: 1053–1054.

94. Egawa, N, Yamamoto, K, Inoue, H, Hikawa, R, Nishi, K, Mori, K, et al. (2011). The endoplasmic reticulum stress sensor, ATF6alpha, protects against neurotoxin-induced dopaminergic neuronal death. J Biol Chem 286: 7947–7957.

95. Hashida, K, Kitao, Y, Sudo, H, Awa, Y, Maeda, S, Mori, K, et al. (2012). ATF6alpha promotes astroglial activation and neuronal survival in a chronic mouse model of Parkinson’s disease. PLoS One 7: e47950.

96. Luk, KC, Kehm, VM, Zhang, B, O’Brien, P, Trojanowski, JQ, and Lee, VM (2012). Intracerebral inoculation of pathological alpha-synuclein initiates a rapidly progressive neurodegenerative alpha-synucleinopathy in mice. J Exp Med 209: 975–986.

97. Cornejo, VH, Pihan, P, Vidal, RL, and Hetz, C (2013). Role of the unfolded protein response in organ physiology: lessons from mouse models. IUBMB life 65: 962–975.

98. Aragon, IV, Barrington, RA, Jackowski, S, Mori, K, and Brewer, JW (2012). The specialized unfolded protein response of B lymphocytes: ATF6alpha-independent development of antibody-secreting B cells. Mol Immunol 51: 347–355.

99. Adachi, Y, Yamamoto, K, Okada, T, Yoshida, H, Harada, A, and Mori, K (2008). ATF6 is a transcription factor specializing in the regulation of quality control proteins in the endoplasmic reticulum. Cell structure and function 33: 75–89.

100. Zhang, K, Wang, S, Malhotra, J, Hassler, JR, Back, SH, Wang, G, et al. (2011). The unfolded protein response transducer IRE1alpha prevents ER stress-induced hepatic steatosis. EMBO J 30: 1357–1375.

101. Wu, J, Rutkowski, DT, Dubois, M, Swathirajan, J, Saunders, T, Wang, J, et al. (2007). ATF6alpha optimizes long-term endoplasmic reticulum function to protect cells from chronic stress. Developmental cell 13: 351–364.

102. Marciniak, SJ (2019). Endoplasmic reticulum stress: a key player in human disease. The FEBS journal 286: 228–231.

103. Urra, H, Dufey, E, Avril, T, Chevet, E, and Hetz, C (2016). Endoplasmic Reticulum Stress and the Hallmarks of Cancer. Trends Cancer 2: 252–262.

104. Nam, SM, and Jeon, YJ (2019). Proteostasis In The Endoplasmic Reticulum: Road to Cure. Cancers (Basel) 11.

105. Gonzalez-Teuber, V, Albert-Gasco, H, Auyeung, VC, Papa, FR, Mallucci, GR, and Hetz, C (2019). Small Molecules to Improve ER Proteostasis in Disease. Trends in pharmacological sciences 40: 684–695.

106. Paxman, R, Plate, L, Blackwood, EA, Glembotski, C, Powers, ET, Wiseman, RL, et al. (2018). Pharmacologic ATF6 activating compounds are metabolically activated to selectively modify endoplasmic reticulum proteins. eLife 7.

107. Grandjean, JMD, Madhavan, A, Cech, L, Seguinot, BO, Paxman, RJ, Smith, E, et al. (2020). Pharmacologic IRE1/XBP1s activation confers targeted ER proteostasis reprogramming. Nature chemical biology 16: 1052–1061.

108. Ogen-Shtern, N, Ben David, T, and Lederkremer, GZ (2016). Protein aggregation and ER stress. Brain Res 1648: 658–666.

109. Hoozemans, JJ, Veerhuis, R, Van Haastert, ES, Rozemuller, JM, Baas, F, Eikelenboom, P, et al. (2005). The unfolded protein response is activated in Alzheimer’s disease. Acta neuropathologica 110: 165–172.

110. Cooper, AA, Gitler, AD, Cashikar, A, Haynes, CM, Hill, KJ, Bhullar, B, et al. (2006). Alpha-synuclein blocks ER-Golgi traffic and Rab1 rescues neuron loss in Parkinson’s models. Science 313: 324–328.

111. Gitler, AD, Bevis, BJ, Shorter, J, Strathearn, KE, Hamamichi, S, Su, LJ, et al. (2008). The Parkinson’s disease protein alpha-synuclein disrupts cellular Rab homeostasis. Proc Natl Acad Sci U S A 105: 145–150.

112. Chung, CY, Khurana, V, Auluck, PK, Tardiff, DF, Mazzulli, JR, Soldner, F, et al. (2013). Identification and rescue of alpha-synuclein toxicity in Parkinson patient-derived neurons. Science 342: 983–987.

113. Bellucci, A, Navarria, L, Zaltieri, M, Falarti, E, Bodei, S, Sigala, S, et al. (2011). Induction of the unfolded protein response by alpha-synuclein in experimental models of Parkinson’s disease. J Neurochem 116: 588–605.

114. Belal, C, Ameli, NJ, El Kommos, A, Bezalel, S, Al’Khafaji, AM, Mughal, MR, et al. (2012). The homocysteine-inducible endoplasmic reticulum (ER) stress protein Herp counteracts mutant alpha-synuclein-induced ER stress via the homeostatic regulation of ER-resident calcium release channel proteins. Hum Mol Genet 21: 963–977.

115. Colla, E, Coune, P, Liu, Y, Pletnikova, O, Troncoso, JC, Iwatsubo, T, et al. (2012). Endoplasmic reticulum stress is important for the manifestations of alpha-synucleinopathy in vivo. J Neurosci 32: 3306–3320.

116. Duennwald, ML, and Lindquist, S (2008). Impaired ERAD and ER stress are early and specific events in polyglutamine toxicity. Genes & development 22: 3308–3319.

117. Kalathur, RK, Giner-Lamia, J, Machado, S, Barata, T, Ayasolla, KR, and Futschik, ME (2015). The unfolded protein response and its potential role in Huntington’s disease elucidated by a systems biology approach. F1000Res 4: 103.

118. Piguet, F, Alves, S, and Cartier, N (2017). Clinical Gene Therapy for Neurodegenerative Diseases: Past, Present, and Future. Hum Gene Ther 28: 988–1003.

119. Hudry, E, and Vandenberghe, LH (2019). Therapeutic AAV Gene Transfer to the Nervous System: A Clinical Reality. Neuron 101: 839–862.

120. Mercado, G, Valdes, P, and Hetz, C (2013). An ERcentric view of Parkinson’s disease. Trends Mol Med 19: 165–175.

121. Michel, PP, Hirsch, EC, and Hunot, S (2016). Understanding Dopaminergic Cell Death Pathways in Parkinson Disease. Neuron 90: 675–691.

122. Hoozemans, JJ, van Haastert, ES, Nijholt, DA, Rozemuller, AJ, and Scheper, W (2012). Activation of the unfolded protein response is an early event in Alzheimer’s and Parkinson’s disease. Neurodegener Dis 10: 212–215.

123. Hoozemans, JJ, van Haastert, ES, Eikelenboom, P, de Vos, RA, Rozemuller, JM, and Scheper, W (2007). Activation of the unfolded protein response in Parkinson’s disease. Biochem Biophys Res Commun 354: 707–711.

124. Mercado, G, Lopez, N, Martinez, A, Sardi, SP, and Hetz, C (2018). A new model to study cell-to-cell transfer of alphaSynuclein in vivo. Biochem Biophys Res Commun 503: 1385–1393.

125. Pobre, KFR, Poet, GJ, and Hendershot, LM (2019). The endoplasmic reticulum (ER) chaperone BiP is a master regulator of ER functions: Getting by with a little help from ERdj friends. J Biol Chem 294: 2098–2108.

126. Salganik, M, Sergeyev, VG, Shinde, V, Meyers, CA, Gorbatyuk, MS, Lin, JH, et al. (2015). The loss of glucose-regulated protein 78 (GRP78) during normal aging or from siRNA knockdown augments human alpha-synuclein (alpha-syn) toxicity to rat nigral neurons. Neurobiol Aging 36: 2213–2223.

127. Gorbatyuk, MS, Knox, T, LaVail, MM, Gorbatyuk, OS, Noorwez, SM, Hauswirth, WW, et al. (2010). Restoration of visual function in P23H rhodopsin transgenic rats by gene delivery of BiP/Grp78. Proc Natl Acad Sci U S A 107: 5961–5966.

128. Vidal, RL, Figueroa, A, Court, FA, Thielen, P, Molina, C, Wirth, C, et al. (2012). Targeting the UPR transcription factor XBP1 protects against Huntington’s disease through the regulation of FoxO1 and autophagy. Hum Mol Genet 21: 2245–2262.

129. Camacho, C, Coulouris, G, Avagyan, V, Ma, N, Papadopoulos, J, Bealer, K, et al. (2009). BLAST+: architecture and applications. BMC Bioinformatics 10: 421.

130. Burley, SK, Berman, HM, Bhikadiya, C, Bi, C, Chen, L, Di Costanzo, L, et al. (2019). RCSB Protein Data Bank: biological macromolecular structures enabling research and education in fundamental biology, biomedicine, biotechnology and energy. Nucleic acids research 47: D464–D474.

131. Madeira, F, Park, YM, Lee, J, Buso, N, Gur, T, Madhusoodanan, N, et al. (2019). The EMBL-EBI search and sequence analysis tools APIs in 2019. Nucleic acids research 47: W636–W641.

132. Hooft, RW, Sander, C, Scharf, M, and Vriend, G (1996). The PDBFINDER database: a summary of PDB, DSSP and HSSP information with added value. Comput Appl Biosci 12: 525–529.

133. Krivov, GG, Shapovalov, MV, and Dunbrack, RL, Jr. (2009). Improved prediction of protein side-chain conformations with SCWRL4. Proteins 77: 778–795.

134. Weirauch, MT, Yang, A, Albu, M, Cote, AG, Montenegro-Montero, A, Drewe, P, et al. (2014). Determination and inference of eukaryotic transcription factor sequence specificity. Cell 158: 1431–1443.

135. Ryno, LM, Genereux, JC, Naito, T, Morimoto, RI, Powers, ET, Shoulders, MD, et al. (2014). Characterizing the altered cellular proteome induced by the stress-independent activation of heat shock factor 1. ACS Chem Biol 9: 1273–1283.

136. Perez-Riverol, Y, Csordas, A, Bai, J, Bernal-Llinares, M, Hewapathirana, S, Kundu, DJ, et al. (2019). The PRIDE database and related tools and resources in 2019: improving support for quantification data. Nucleic acids research 47: D442–D450.

137. O’Riordan, CR, Lachapelle, AL, Vincent, KA, and Wadsworth, SC. Scaleable chromatographic purification process for recombinant adeno-associated virus (rAAV). The journal of gene medicine 2: 444–454.

138. Mahul-Mellier, AL, Vercruysse, F, Maco, B, Ait-Bouziad, N, De Roo, M, Muller, D, et al. (2015). Fibril growth and seeding capacity play key roles in alpha-synuclein-mediated apoptotic cell death. Cell Death Differ 22: 2107–2122.

